# NCOR2 represses MHC class I molecule expression to drive metastatic progression of breast cancer

**DOI:** 10.1101/2025.03.10.642060

**Authors:** Pavla Ticha, Jason J. Northey, Kelly Kersten, Hugo González Velozo, Alastair J. Ironside, Martin Zidek, Allison Drain, Jonathan N. Lakins, Yunn-Yi Chen, Kelvin K. Tsai, Valerie M. Weaver

## Abstract

Metastatic progression depends upon the ability of disseminated tumor cells to evade immune surveillance. MHC molecule expression facilitates T cell recognition and activation to permit the eradication of metastatic tumor cells. We identified nuclear corepressor 2 (NCOR2) as a key epigenetic regulator of MHC class I molecule expression on breast tumor cells. Patients with triple negative breast cancers (TNBC) that expressed high levels of NCOR2 also exhibited reduced metastasis free survival and decreased MHC class I expression, and the metastatic lesions in patients with TNBC had high nuclear NCOR2 and reduced CD8 T cell levels and activity. Genetically and experimentally reducing NCOR2 expression in tumor cells permitted interferon gamma upregulation of MHC class I, and potentiated CD8 T cell activity and induction of apoptosis to repress metastatic progression of disseminated breast cancer cells. These studies provide evidence to support NCOR2 as a targetable epigenetic regulator of metastasis towards which therapies could be developed to reduce patient mortality.

## INTRODUCTION

Cancer patient mortality is primarily due to treatment resistant, metastatic disease (1). Tumor cells can disseminate early and either remain dormant for years or generate progressive, metastatic disease (2–8). Metastatic progression versus dormancy depend upon the growth and survival of the disseminated tumor cells (DTCs) at the metastatic site, (9–14) as well as the ability of the DTCs to evade immune surveillance (15–18). Of these determinants, intrinsic tumor cell and extrinsic systemic and microenvironmental factors that regulate anti-tumor immunity have emerged as key regulators of tumor dormancy versus progression (2, 3, 9, 16, 19–30). Clarifying mechanisms whereby disseminated tumor cells overcome antitumor immunity to generate progressive metastatic lesions is an area of intense investigation.

The expression of MHC molecules on the surface of cancer cells and their tumor peptide presentation facilitates T cell recognition and the eradication of DTCs (31–34). The amount and type of MHC class molecule expressed on tumor cells also influences CD4 and CD8 T cell activity to amplify their antitumor activity (35–39). Consistently, loss of MHC molecule cell surface expression has been implicated in progressive metastatic disease (31). Mechanisms regulating dysfunctional levels and function of MHC molecules in tumors include irreversible genetic mutations or polymorphisms in the MHC molecule or its accessory proteins (40). These irreversible genetic perturbations include downregulation or mutations/deletions in Human Leukocyte Antigens (HLA; human MHC molecules) alleles that reduce T cell recognition (41–43) or accessory molecules that influence cell surface expression (44), as well as polymorphisms that compromise tumor peptide binding and presentation (45, 46). By contrast epigenetic silencing of MHC or associated processing molecules, modification of pathways that regulate transcription of MHC or its accessory molecules including NFκB or interferon gamma (IFNγ)(47), or factors influencing post translational processing, cell surface expression and protein stability (48–51), represent reversible mechanisms that could be targeted to potentiate antitumor immunity and eradicate metastatic burden.

Women with triple negative breast cancer (TNBC) and Herceptin receptor positive (HER2+ve) breast cancers exhibit high rates of progressive metastatic disease, despite successful primary tumor removal and comprehensive chemotherapy (52, 53). Developing targeted therapies to eradicate disseminated tumor cells with high potential to form metastatic lesions would go far towards enhancing breast cancer patient survival. We identified nuclear corepressor 2 (NCOR2) as an epigenetic regulator of treatment resistance in TNBC (54). We and others showed that patients whose breast cancers express high levels of NCOR2 also exhibit overall poor prognosis and reduced survival (54, 55). We showed that NCOR2 represses apoptosis induction in response to chemotherapy and IFNγ, and that NCOR2 compromises immune checkpoint response by recruiting HDAC3 to repress the expression of pro-apoptotic genes and cytokines implicated in CD8 T cell recruitment (54). Given associations between treatment resistance, metastatic disease and immune evasion, and data indicating IFNγ induces MHC molecule expression, here we asked whether NCOR2 could regulate tumor metastasis by epigenetically repressing IFNγ-dependent MHC class molecule expression.

## RESULTS

### Nuclear NCOR2 is enriched in residual chemotherapy-treated human breast tumors

Approximately seventy percent of TNBCs that develop in patients exhibit a partial response to neoadjuvant chemotherapy and retain some level of residual disease (56). We previously implicated NCOR2 in systemic chemotherapy resistance in experimental and clinical TNBC (57). Accordingly, here we asked whether NCOR2 expression in the residual TNBC tumors could explain their treatment resistance. Immunohistochemistry analysis of the residual TNBCs excised from a cohort of patient biospecimens that had received neoadjuvant chemotherapy showed elevated nuclear NCOR2 staining when compared to an independent cohort of nontreated TNBC tumors (Figure 1A and 1B). The findings are consistent with the possibility that NCOR2 contributed to the treatment resistance phenotype of these tumors.

**Fig. 1.**
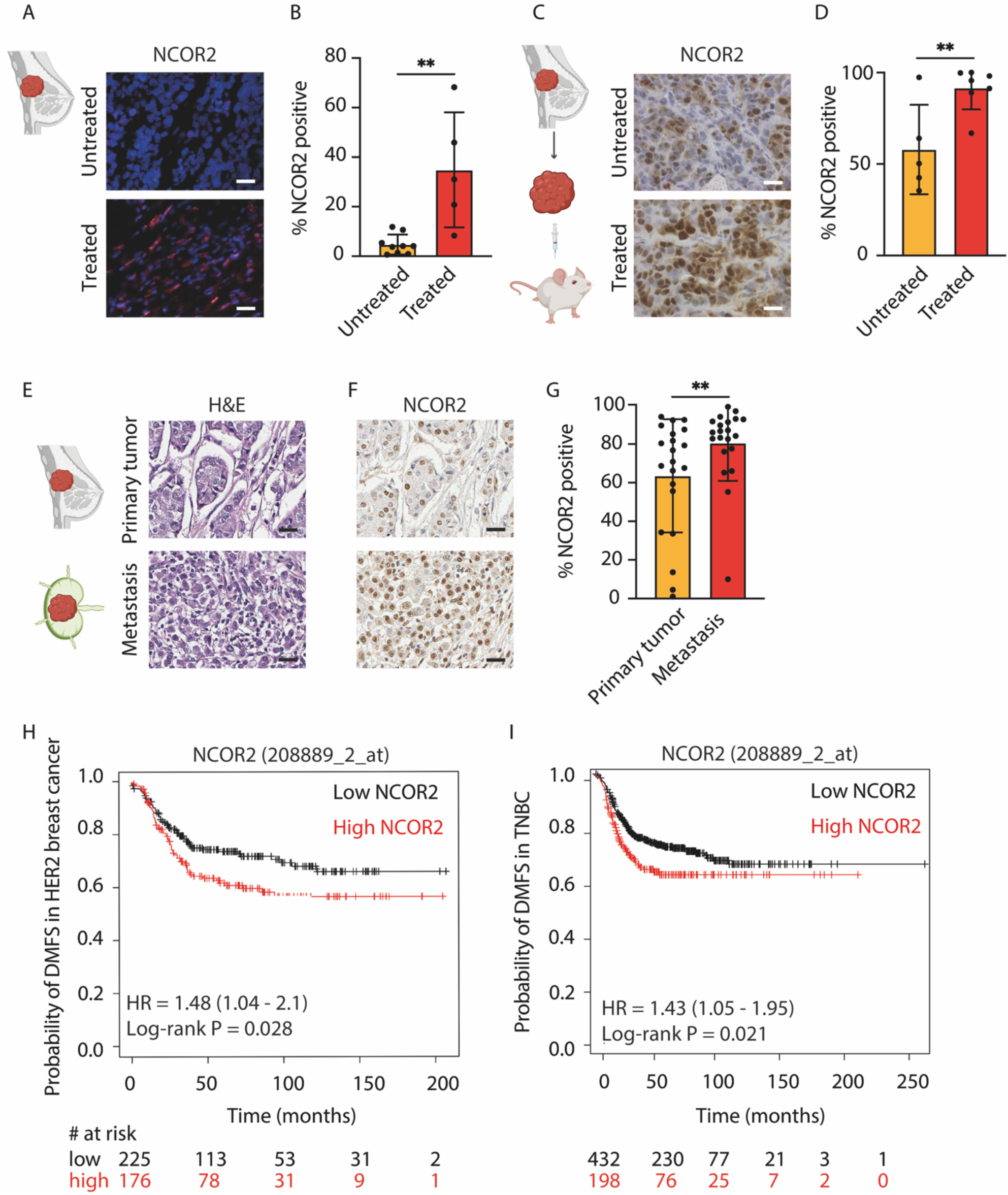
NCOR2 is enriched in residual chemotherapy treated human breast tumors and predicts reduced metastasis-free survival. (**A**) Representative images of frozen tissue sections from systemic chemotherapy treated and untreated human triple-negative breast cancer (TNBC) stained for NCOR2. Scale bar: 20 μm. (**B**) Bar graphs showing quantification of positive IHC NCOR2 staining shown in (A) (untreated, *n* = 9; treated, *n* = 5). (**C**) Representative images of paraffin sections from systemic chemotherapy treated and untreated TNBC patient-derived xenograft (PDX) primary tumors stained for NCOR2. Scale bar: 20 μm. (**D**) Bar graphs showing quantification of positive immunohistochemical (IHC) NCOR2 staining shown in (C) (untreated, *n* = 5; treated, *n* = 7). (**E**) Representative images of paraffin sections from human primary ductal carcinoma and matched lymph node metastasis stained for hematoxylin and eosin (H&E). Scale bar: 20 μm. (**F**) Representative images of IHC staining for NCOR2 of paraffin sections described in (E). Scale bar: 20 μm. (**G**) Bar graphs showing quantification of positive IHC staining for NCOR2 shown in (F) (primary tumor, *n* = 23; metastasis, *n* = 23). (**H**) Kaplan-Meier plot (using http://kmplot.com) of the distant metastasis-free survival (DMFS) of HER-2 positive breast cancer patients split by NCOR2 mRNA abundance (high or low) in primary tumors (*n* = 451; P = 0.028). The hazard ratio (HR) is shown. (**I**) Kaplan-Meier plot (analyzed using http://kmplot.com) of the DMFS of basal breast cancer patients split by NCOR2 mRNA abundance (high or low) in primary tumors (*n* = 630; P = 0.021). The hazard ratio (HR) is shown. Graphs show mean ± s.e.m. *P < 0.05, **P < 0.01 (two-tailed Mann-Whitney *U* test).

To test whether chemotherapy treatment led to the enrichment of NCOR2 in therapy resistant human breast tumors, we next treated human patient derived xenograft (PDX) TNBCs propagated orthotopically in NOD/SCID mice with Paclitaxel (10 mg/kg for 4 weeks, every second day starting 7 days after PDX implantation) and thereafter stained the treated and untreated tissues for NCOR2 protein. We quantified a significant increase in immunohistochemical staining for nuclear NCOR2 in the residual chemotherapy-treated resistant TNBC PDX tumors when compared to the cytosolic and heterogeneous nuclear levels of NCOR2 in the non-treated tumors (Figure 2C and 2D). The data implicate NCOR2 in the residual treatment resistant human TNBC phenotype.

**Fig. 2.**
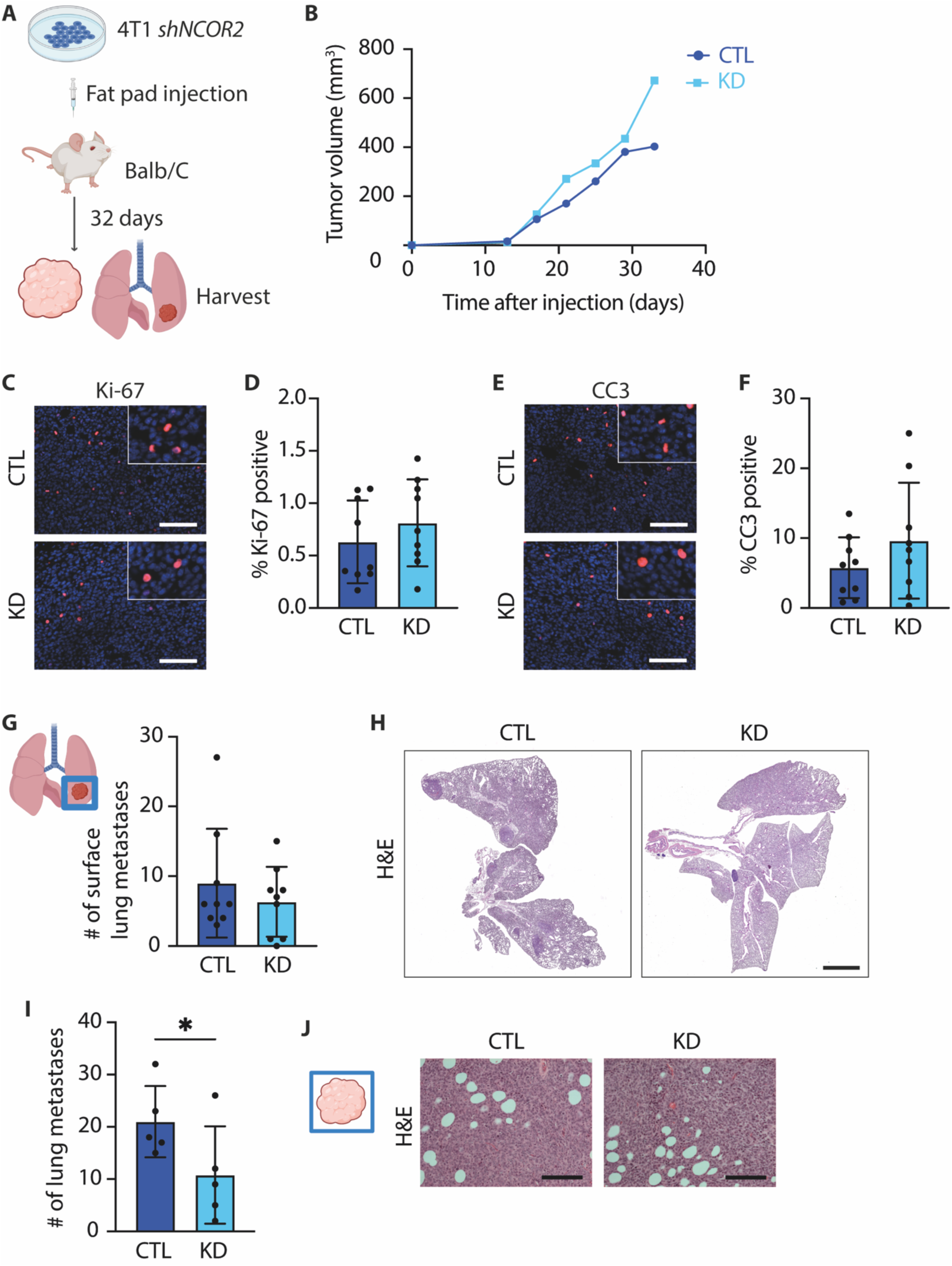
NCOR2 regulates lung metastasis from primary syngeneic mammary tumors. (**A**) Cartoon showing experimental design. **(B**) Line graph showing rate of tumor volume increase in 4T1 tumor cells injected into Balb/C mice with (KD) and without (CTL) NCOR2 knockdown. (**C**) Representative images of IHC staining for Ki-67 in sections from tissue as described in (B). Scale bar: 100 μm. (**D**) Bar graphs showing quantification of IHC staining for Ki-67 staining as shown in (C) (control, *n* = 9; NCOR2 knockdown, *n* = 9). (**E**) Representative images of IHC of cleaved caspase-3 (CC3) stained sections from tissue described in (B). Scale bar: 100 μm. (**F**) Bar graphs showing quantification of IHC Ki-67 staining shown in (E) (control, *n* = 9; NCOR2 knockdown, *n* = 9). (**G**) Bar graph showing quantification of the number of surface lung metastases from specimens described in (A) (control, *n* = 9; NCOR2 knockdown, *n* = 9). (**H**) Representative images of paraffin sections of lungs from Balb/C mice orthotopically injected with 4T1 cells with (KD) or without (CTL) NCOR2 knockdown and stained for H&E. Scale bar: 1 mm. (**I**) Bar graphs showing quantification of the number of lung metastases in specimens described in (H) (control, *n* = 9; NCOR2 knockdown, *n* = 9). (**J**) Representative images of paraffin sections from 4T1 syngeneic primary tumors stained by H&E. Scale bar: 100 μm. Graphs show mean ± s.e.m. *P < 0.05 (one-tailed Mann-Whitney *U* test).

### Metastatic human breast cancer cells are enriched in nuclear NCOR2 and high NCOR2 predicts lower metastasis-free survival in breast cancer patients

An incomplete pathological response to therapy associates with a worse overall survival in patients with TNBC (56). TNBCs that fail to respond with a complete pathological response to chemotherapy frequently recur with metastatic disease that is refractory to multiple systemic therapies, with less than 30% of patients surviving beyond 5 years (58). To determine whether there were links between NCOR2 and human breast cancer metastasis, we quantified the level of nuclear NCOR2 in tissue arrayed sections of human primary invasive ductal carcinoma and compared to patient matched lymph node metastasis. Quantitative immunohistochemical analysis revealed that there was a significant enrichment for nuclear NCOR2 in the tumor cells that had disseminated to the human lymph nodes in patients with TNBC, as compared to the levels found in their patient matched primary invasive TNBC (Figure 1E-G). In further support of an association between NCOR2 and reduced overall patient survival, the distant metastasis-free survival (DMFS) of basal breast cancer patients split by NCOR2 mRNA abundance (high or low) showed that patients with high NCOR2 in their primary tumors had a less favorable DMFS (Figure 1H). Also for HER-2 positive breast cancer patients split by NCOR2 mRNA abundance (high or low), patients with high NCOR2 in their primary tumors similarly had a less favorable DMFS (Figure 1I). These findings implicate NCOR2 in breast cancer metastasis and extend the observations beyond TNBCs.

### NCOR2 regulates syngeneic mammary tumor lung metastasis

We next examined the role of NCOR2 in experimental breast cancer metastasis. We used the highly invasive and metastatic TNBC murine line 4T1, derived from a tumor that developed within a Balb/C mouse that spontaneously forms progressive metastatic lesions in the liver, lung, brain and bone of host syngeneic mice (59, 60). We engineered efficient reduction in NCOR2 protein levels in the 4T1 cells using a construct designed for Isopropyl ß-D-1-thiogalactopyranoside (IPTG)-inducible expression of NCOR2-targeting shRNA (KD) or a GFP-targeting control shRNA (CTL) (Extended Data Figure 1A and 1B). Thereafter, we orthtopically injected the engeneered cells with and without NCOR2 knockdown into syngeneic hosts and confrimed that the shRNA-mediated reduction in NCOR2 levels were maintained in mammary tumors *in vivo* (Figure 2A and Extended Data Figure 1C). Tumor volume measurements throughout tumor progression suggested that NCOR2 expression had little to no impact on primary tumor growth (Figure 2B). Immunohistochemistry of sections from tumor tissue sections excised at study termination revealed similar levels of proliferation and apoptosis, as indicated by staining for Ki-67 and cleaved (activated) caspase 3 (Figure 2C-F). Importantly, despite a lack of differences in primary tumor growth and survival, macroscopic examination of the lungs revealed there was a significantly reduced number of lung surface metastases in the NCOR2 knockdown cohort as compared to the control group (Figure 2G). Hematoxylin and eosin (H&E) staining of the lung tissues from these mice further demonstrated that there was a lower number of macrometastases in the 4T1 tumors in which NCOR2 was knocked down as compared to the control 4T1 injected tumor group (Figure 2H). Further analysis revealed that the metastatic lung lesions in the mice in which NCOR2 was knocked down were smaller than those that formed in the mice injected with 4T1 control tumor cells (Figure 2I). H&E staining of the primary tumor tissue showed similar morphology and histophenotype between the tumors with and without NCOR2 knockdown (Figure 2J), implying NCOR2 likely tempered tumor metastasis either by influencing tumor cell dissemination or the metastatic outgrowth of the tumor cells in the lungs.

### NCOR2 regulates lung metastasis from spontaneous mammary tumors

To more rigorously explore the role of NCOR2 in breast cancer progression we generated cohorts of FVB MMTV-PyMT; MMTV-Cre; NCOR2^+/+^, NCOR2^-/+^ and NCOR2^-/-^ mice to assess the impact of NCOR2 on spontaneous tumor behavior (Extended Data Figure 2A), and confirmed genomic deletion and mammary tumor specific NCOR2 knockout in the experimental group (Extended Data Figure 2B-E). The PyMT NCOR2 mouse cohorts were then monitored for differences in spontaneous tumor formation and phenotype, circulating tumor cell (CTC) levels and lung metastasis. Primary tumor latency was similar in all three mouse cohorts (Figure 3A), as was tumor volume as assessed by weekly caliper measurements (Figure 3B). Mouse survival, as indicated by time when primary tumor size reached a size of 1.5 cm in diameter (Figure 3C), and the number of tumors per mouse also did not differ between the experimental groups (Figure 3D). Pathological examination of H&E stained primary breast tumor tissue (Figure 3E) indicated that the tumors formed in all three of the mouse cohorts were similar in histophenotype. In each cohort the tumors were moderately or poorly differentiated adenocarcinoma with pappillary, trabecular and acinar growth patterns showing varying degrees of differentiation (nuclei showed increased amount of mitotic activity and pleomorphism) with about a third of the tumors showing significant necrosis (>10%). Immunohistochemical analysis of primary excised tumors confirmed there were no discernible differences in proliferation or apoptosis between the three tumor genotypes, as indicated by Ki-67 (Figure 3F and 3G), and cleaved Caspase 3 (CC3) staining (Figure 3H and 3I). These findings indicated that NCOR2 expression had little to no impact on the primary tumor histophenotype, growth and apoptosis of endogenous mammary tumors. However, similar to the results obtained in the 4T1 syngeneic studies, macroscopic examination of lungs excised at study termination showed a significantly reduced number of surface metastatic nodules in the PyMT NCOR2^-/+^ and PyMT NCOR2^-/-^ mice as compared to the PyMT NCOR2^+/+^ mice (Figure 3J). Further histological examination suggested there was a significantly lower metastatic burden in the PyMT NCOR2^-/+^ or PyMT NCOR2^-/-^ mice relative to the wild type PyMT-NCOR2^+/+^ mice (Figure 3K). H&E stained lung tissues showed a significantly higher number of macrometastases in the PyMT NCOR2^+/+^ mice that quantitative analysis revealed were also significantly larger (Figure 3L-N). Importantly, CTC levels were not significantly different between the three experimental mouse cohorts suggesting NCOR2 had little to no impact on metastatic dissemination (Figure 3O). Instead, the findings suggest that NCOR2 influences the metastatic outgrowth of DTCs after they have reached their distal site. Consistent with this idea, when we combined the total number of micro and macro metastases, we detected no significant difference in the total number of metastatic lesions between the three experimental groups (Figure 3P). Interestingly, although we quantified no differences in the proliferation of the metastatic lesions (Figure 3Q and 3R) we did observe a significant increase in apoptosis in the lung disseminated lesions in the NCOR2 heterozygous mice (Figure 3S and 3T). These findings confirm that although NCOR2 has little impact on primary tumor development and growth it significantly affects the metastatic survival of disseminated tumor cells to regulate their metastatic outgrowth.

**Fig. 3.**
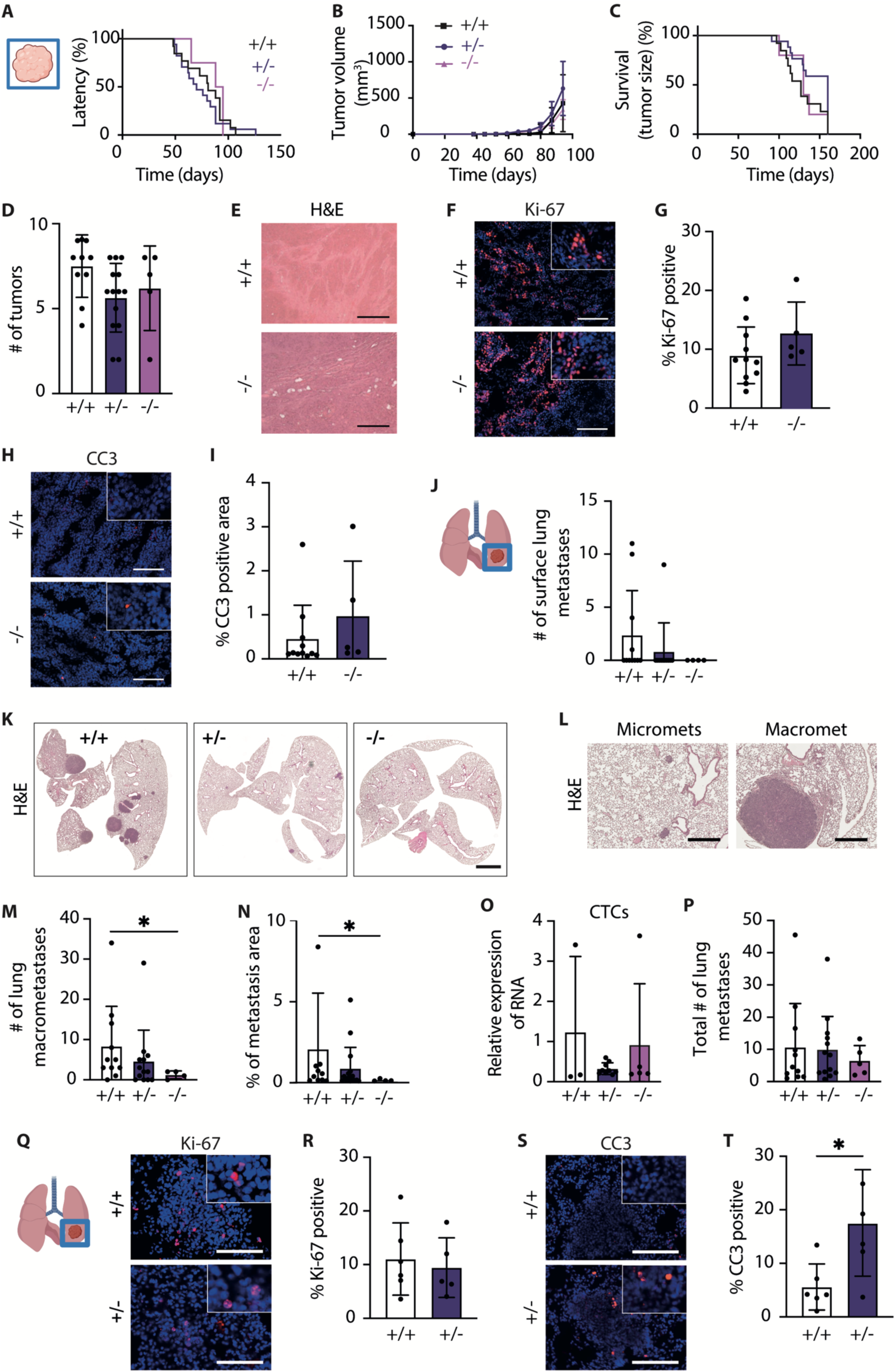
NCOR2 regulates lung metastasis in PyMT-NCOR2 knockout mice. (**A**) Line graph showing tumor latency (days) in the PyMT wildtype (+/+), PyMT MMTV-Cre NCOR2^+/-^, and PyMT MMTV-Cre NCOR2^-/-^ transgenic mice (PyMT wildtype, *n* = 10; PyMT NCOR2^+/-^, *n* = 14; PyMT NCOR2^-/-^; *n* = 5. (**B**) Line graph showing increase in tumor volume in each of the three transgenic groups from (A). (**C**) Line graph illustrating mouse survival plots for each transgenic cohort from (A). (**D**) Bar graphs showing quantification of primary tumor number per mouse in each of the transgenic mouse cohorts (PyMT wildtype, *n* = 10; PyMT NCOR2^+/-^, *n* = 14; PyMT NCOR2^-/-^; *n* = 5). (**E**) Representative images of H&E-stained paraffin sections from the PyMT wildtype (+/+) and PyMT NCOR2^-/-^ transgenic mice. Scale bar: 100 μm. (**F**) Representative images of IHC staining for Ki-67 in frozen sections from each of the primary transgenic tumors. Scale bar: 100 μm. (**G**) Bar graphs showing quantification of positive IHC staining shown in (F) (PyMT wild type, *n* = 11; PyMT NCOR2^-/-^, *n* = 5). (**H**) Representative IHC images of frozen sections of tumors as in (F) stained for Cleaved Caspase 3 (CC3). Scale bar: 100 μm. **(I)** Bar graphs showing quantification of IHC stained tissue shown in (H) (PyMT wild type, *n* = 11; PyMT NCOR2^-/-^, *n* = 5). (**J**) Bar graphs showing quantification of the number of surface lung metastases for mouse cohorts from (A) (PyMT wild type, *n* = 11; PyMT NCOR2^+/-^, *n* = 13; PyMT NCOR2^-/-^, *n* = 4). **(K)** Representative images of paraffin sections of lungs stained for H&E. Scale bar: 1 mm. (**L**) Representative images of paraffin sections from lung parenchyma with micrometastases or macrometastasis stained for H&E. Scale bar: 200 μm. (**M**) Bar graphs showing quantification of the number of lung macrometastases shown in (K) (PyMT wild type, *n* = 11; PyMT NCOR2^+/-^, *n* = 13; PyMT NCOR2^-/-^, *n* = 4). (**N**) Bar graph quantifying the percentage of metastasis area in the lungs for each of the groups shown in (K) (PyMT wild type, *n* = 11; PyMT NCOR2^+/-^, *n* = 13; NCOR2^-/-^, *n* = 4). (**O**) Bar graphs showing quantification of the relative levels of CTCs (circulating tumor cells) as assessed by RT-qPCR for PyMT in each of the murine cohorts (PyMT wild type, *n* = 3; PyMT NCOR2^+/-^, *n* = 9; PyMT NCOR2^-/-^, *n* = 4). (**P**) Bar graphs showing quantification of the total number of lung metastases shown in (K) (PyMT wild type, *n* = 11; PyMT NCOR2^+/-^, *n* = 13; PyMT NCOR2^-/-^, *n* = 4). (**Q**) Representative images showing IHC staining for Ki-67 in paraffin sections from lung tissue excised from PyMT wild type (+/+) and PyMT NCOR2^+/-^ mice. Scale bar: 100 μm. (**R**) Bar graphs showing quantification of IHC staining shown in (Q) (PyMT wild type, *n* = 6; PyMT NCOR2^+/-^, *n* = 5). (**S**) Representative images showing IHC staining for CC3 in paraffin sections of lungs excised from mouse cohorts as in (Q). Scale bar: 100 μm. (**T**) Bar graphs showing quantification of IHC staining shown in (S) (PyMT wild type, *n* = 6; PyMT NCOR2^+/-^, *n* = 5). Graphs show mean ± s.e.m. *P < 0.05 (two-tailed Mann-Whitney *U* test).

### NCOR2 regulates metastatic outgrowth of breast cancer cells

To explore the possibility that NCOR2 directly regulates the survival of disseminated tumor cells to modulate their metastatic outgrowth we exploited the less aggressive murine 4T07 breast cancer cell line. 4T07 breast cancer cells are unable to spontaneously form lung metastasis when injected into the mammary fat pad of host syngeneic mice, however these tumor cells will form metastatic lesions in the lungs when they are injected via tail vein. (59) After confirming that the 4T07 breast cancer cells expressed abundant levels of NCOR2 protein, we transduced them with the same lentiviral shRNA constructs used to knock down NCOR2 (KD, and GFP control; CTL) in the 4T1 cells and verified efficient reduction of NCOR2 protein expression that was maintained in mammary tumors *in vivo* (Extended Data Figure 3A-C). We then injected the tumor cells into the tail vein of host syngeneic mice and monitored them twice per week for metastatic lung growth using bioluminescent imaging (BLI) (Figure 4A). Two weeks following tail vein injection we detected a strong BLI signal in the lungs of the mice that were injected with the 4T07 cells expressing the control vector, and by comparison a substantially weaker signal in the mice that had been injected with the cells in which NCOR2 had been knocked down (Figure 4B and 4C). Consistently, at study termination, H&E staining revealed a strikingly lower metastatic burden in the lungs of the mice that had been injected with the 4T07 cells in which NCOR2 had been knocked down as compared to the cells that expressed the control shRNA (Figure 4D). Further analysis revealed that there was a significantly reduced number of macrometastases and overall decreased metastatic lesion area using the same comparison (Figure 4E and 4F). Immunofluorescence staining for Ki-67 revealed similar levels of proliferation in the lung metastases in both experimental groups (Figure 4G and 4H). However, apoptosis levels in the macro-metastatic lung lesions that developed in the mice harboring the 4T07 breast cancer cells in which NCOR2 was knocked down were significantly higher, as indicated by more than a three fold increase in detectable CC3 staining (Figure 4I and 4J). These findings demonstrate that NCOR2 regulates metastatic outgrowth in the lung, and implicate reduced tumor cell survival as a plausible mechanism regulating this behavior.

**Fig. 4.**
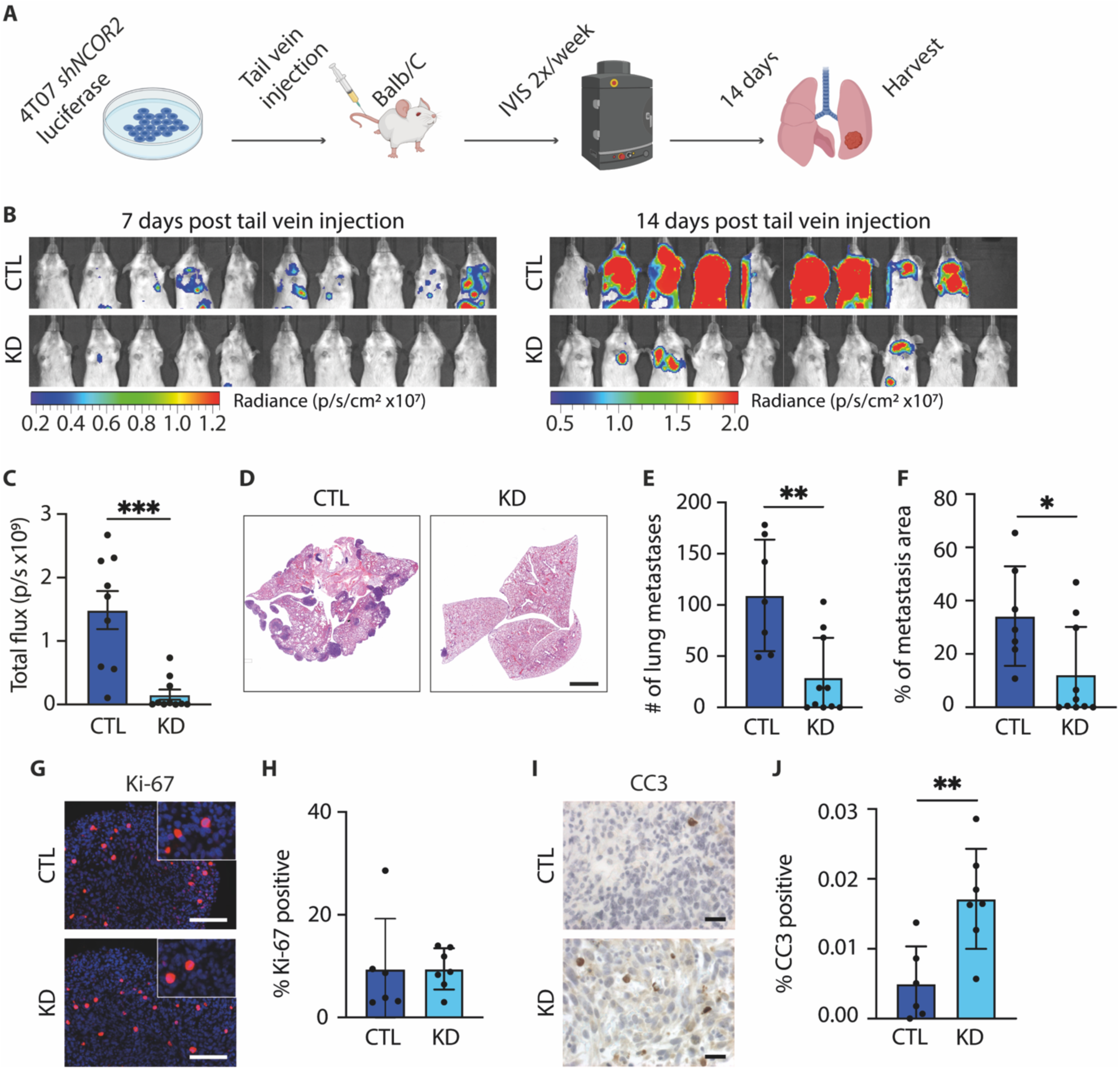
NCOR2 regulates lung metastatic outgrowth following tail vein injection. (**A**) Cartoon showing experimental design. (**B**) Representative bioluminescence images of Balb/C mice 7 days and 14 days after tail vein injection with 4T07 cells with (KD) or without (CTL) knockdown of NCOR2. (**C**) Quantification of Bioluminescence (BLI; total flux) of mouse lung regions from the 14 day time point in (B) (control, *n* = 9; 4T07 NCOR2 knockdown *n* = 10). **(D)** Representative images of paraffin sections of lungs harvested 14 days after tail vein injection, stained for H&E. Scale bar: 1 mm. (**E**) Bar graphs showing quantification of the number of lung metastases shown in (D) (control, *n* = 8; 4T07 NCOR2 knockdown *n* = 10). (**F**) Bar graphs showing quantification of the percent of metastasis area in the lungs shown in (D) (4T07 control, *n* = 8; NCOR2 knockdown, *n* = 10). (**G**) Representative IHC staining for Ki-67 in paraffin sections from lung metastases shown in (D). Scale bar: 100 μm. (**H**) Bar graphs showing quantification of IHC staining shown in (G) (control, *n* = 7; 4T07 NCOR2 knockdown, *n* = 7). (**I**) Representative images of IHC staining for CC3 in paraffin sections of lung metastatic tissue as in (G). Scale bar: 20 μm. (**J**) Bar graph quantifying the IHC staining shown in (I) (control, *n* = 7; 4 NCOR2 knockdown, *n* = 7). Graphs show mean ± s.e.m. *P < 0.05, **P < 0.01 ***P < 0.01 (unpaired t-test).

### NCOR2-dependent metastatic outgrowth depends upon a competent immune system

Metastatic outgrowth is curtailed by anti-tumor immunity (61). We previously demonstrated that NCOR2 represses the expression of multiple cytokines that regulate inflammation and stimulate CD8 T cell recruitment (57). To test whether NCOR2 permits metastatic outgrowth by modulating anti-tumor immunity, we repeated the 4T07 NCOR2 knockdown tail vein studies in a cohort of NOD SCID gamma (NSG) severely immunodeficient mice (62). The 4T07 cells desscribed above (Figure 4) were injected via tail vein into a cohort of NSG mice which were then monitored for metastatic lung growth using bioluminescent imaging (Figure 5A). Although the bioluminescent signal in the lungs of the NSG mice initially indicated a modest reduction in signal in the tumor cells in which NCOR2 was knocked down as compared to CTL cells (Figure 5B), we detected no significant differences in metastatic burden by bioluminescence signal between the two groups at experimental endpoint (Figure 3B and 3C). Consistently, H&E staining of excised lungs revealed a similar level of metastatic burden between the two experimental groups (Figure 5E), including near identical visible macrometastases and percent metastatic area within the whole lung (Figure 5F and 5G). Moreover, immunohistochemical staining for Ki-67 and cleaved (active) caspase-3 in the lungs of the host mice revealed similar levels of proliferation and apoptosis between the two groups (Figure 5H-K). These findings imply NCOR2 influences the metastatic outgrowth of breast cancer cells in the lungs by modulating anti-tumor immunity.

**Fig. 5.**
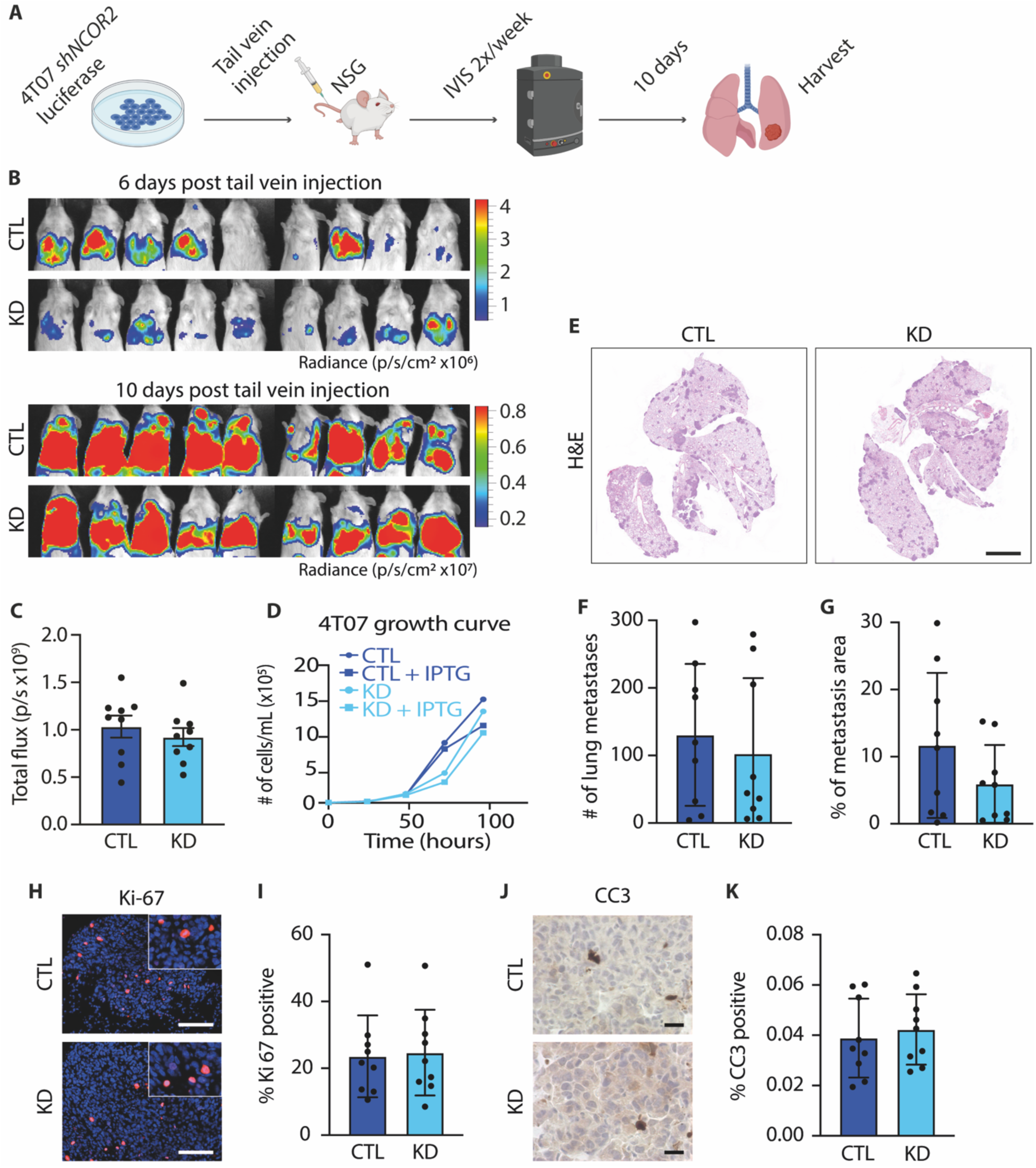
NCOR2 does not regulate lung metastasis in 4T07 tail vein injected immunodeficient mice. (**A**) Cartoon showing experimental design. (**B**) Representative bioluminescence images of NSG mice 6 and 10 days following tail vein injection with 4T07 cells with (KD) or without (CTL) knockdown of NCOR2. (**C**) Quantification of Bioluminescence (BLI; total flux) of mouse lung regions from the 10 day time point in (B) (control, *n* = 9; 4T07 NCOR2 knockdown *n* = 9). (**D**) Line graph showing growth curve of cultured 4T07 cells with or without knockdown of NCOR2. (**E**) Representative images of paraffin sections of lungs stained for H&E. Scale bar: 1 mm. (**F**) Bar graphs showing quantification of the number of lung metastases shown in (E) (control, *n* = 9; NCOR2 knockdown; *n* = 9). (**G**) Bar graphs showing quantification of the percent of metastasis area in the lungs shown in (E) (control, *n* = 9; NCOR2 knockdown; *n* = 9). (**H**) Representative images of IHC staining for Ki-67 in paraffin sections from lung metastases shown in (E). Scale bar: 100 μm. (**I**) Bar graphs showing quantification of IHC staining shown in (H) (control, *n* = 9; NCOR2 knockdown; *n* = 9). (**J**) Representative images of IHC staining for CC3 in paraffin sections from lungs shown in (E). Scale bar: 20 μm. (**K**) Bar graphs showing quantification of IHC staining shown in (J) (control, *n* = 9; NCOR2 knockdown; *n* = 9). Graphs show mean ± s.e.m.

### NCOR2 modulates activity of infiltrating CD8 T cells within metastatic lungs

We next sought to investigate whether NCOR2 regulated the lung metastatic outgrowth of breast cancer cells by modulating CD8 T cell recruitment and/or activity. Flow cytometry analysis of the lungs of mice that had been tail vein injected with 4T07 breast cancer cells with and without NCOR2 knocked down as in Figure 4 revealed a higher overall number of leukocytes (CD45+ cells) in the lungs of the mice injected with breast cancer cells expressing high levels of NCOR2 (Extended Data Figure 4A and 4B). However, a higher percentage of the CD45 cells in the lungs of the mice that had been injected with 4T07 cells in which NCOR2 was knocked down were CD4 and CD8 T cells (Figure 6A-C). Among CD4 T cells, there was a modest increase in the proportion of positive cells and absolute levels (gMFI) of expression of the surface activation marker CD44 (Figure 5D-F). Interestingly, we detected a significant increase in the proportion of positive cells and absolute levels (gMFI) of CD44 on CD8 T cells isolated from the lungs of the mice injected with the 4T07 cells in which NCOR2 was knocked down, indicative of higher levels of activation and enhanced T cell killing potential (Figure 6G-I). Activated CD8 T cells secrete abundant IFNγ that induces cell death in tumor cells (63), and we previously showed that NCOR2 regulates apoptosis sensitivity to IFNγ (57). Consistently, we engineered 4T07 cells to express a second independent NCOR2-targeting shRNA and validated that they produced the same reduced capacity to form progressive metastasis when compared to CTL 4TO7 cells (Extended Data Figure 5). We then treated these same control and NCOR2 knockdown cells with IFNγ in culture and observed the induction of significantly more apoptosis in NCOR2 knockdown cells versus cells expressing high NCOR2 levels (Figure 6J). These findings are consistent with our earlier results showing elevated cell death, indicated by higher activated caspase 3, in the lung metastatic lesions from the spontanenous PyMT tumors in which NCOR2 was genetically knocked out (Figure 3S and 3T), and in the lung metastases present in the Balb/C mice in which the 4T07 cells with NCOR2 knocked down were tail vein injected (Figure 4I and 4J).

**Fig. 6.**
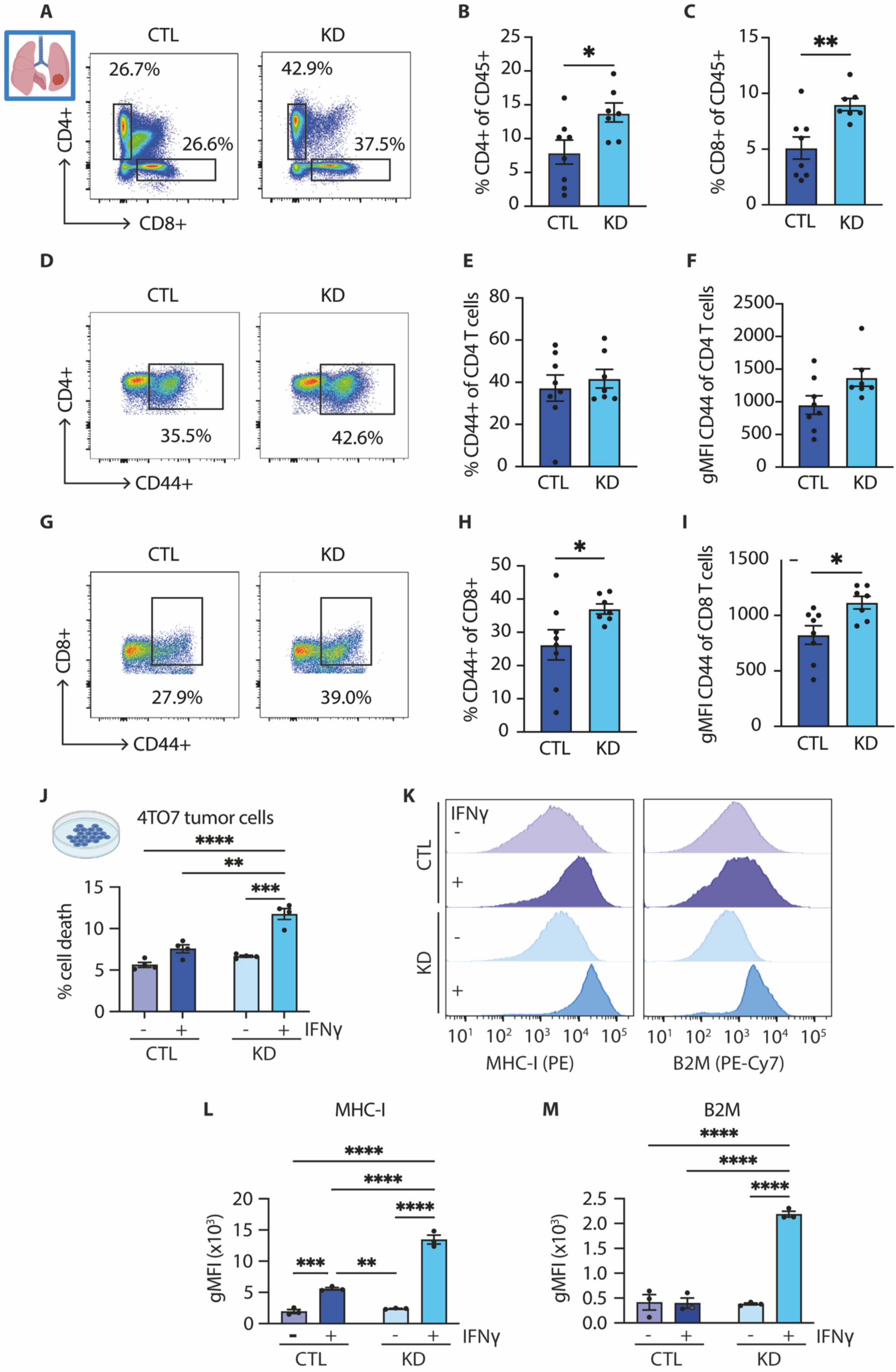
NCOR2 modulates CD8 T cell lung infiltration in 4T07 tail vein injected syngeneic mice. **(A)** Representative flow plots showing the fraction of CD4+ T cells and CD8+ T cells isolated from the lungs of the syngeneic mice intravenously inoculated with 4TO7 cells with (KD) or without (CTL) knockdown of NCOR2 as in Figure 4. **(B)** Bar graphs showing quantification of the percentage of CD4+ T cells, calculated from total CD45+ cells (control, *n* = 8; NCOR2 knockdown, *n* = 7). **(C)** Bar graphs showing quantification of percentage of CD8+ T cells calculated from total CD45+ cells (control, *n* = 8; NCOR2 knockdown, *n* = 7). (**D**) Representative flow plots showing the percentage of CD44+ cells within the CD4+ T cells isolated from the lungs of the syngeneic mice as in (A). (**E**) Bar graphs showing quantification of flow data for (D) (control, *n* = 8; NCOR2 knockdown, *n* = 7). **(F)** Bar graphs showing quantification of gometric Mean Fluorescence Intensity (gMFI) of CD44 surface levels for CD4+ T cells isolated from the lungs of the syngeneic mice shown in (D). (**G**) Representative flow plots showing the percentage of CD44+ cells within the total CD8+ T cells isolated from the lungs of the syngeneic mice as in (A). (**H**) Bar graphs showing quantification of CD44+ CD8+ T cells isolated from the lungs of the syngeneic mice shown in (G) (control, *n* = 8; NCOR2 knockdown, *n* = 7). **(I)** Bar graphs showing quantification of gMFI of CD44 surface levels for CD8+ T cells isolated from the lungs of the syngeneic mice shown in (G). **(J)** Percentage of cell death in 4TO7 with (KD) or without (CTL) knockdown of NCOR2 and stimulated with vehicle or IFNg for 48 hrs (n=3 biological replicates). **(K)** Representative histograms showing cell surface expression for MHC-I and B2M for 4TO7 cells treated as in (J). **(L and M)** Bar graphs showing average gMFI for cell surface levels of MHC-I and B2M for 4TO7 cells treated as in (J) (n=3 biological replicates). Graphs show mean ± s.e.m. *P < 0.05, **P < 0.01 (B-C, E-F, H-I; two-tailed Mann-Whitney *U* test, J, L-M; two-way ANOVA with Tukey’s multiple comparisons test).

Immune cell secreted IFNγ increases MHC class I molecule expression on the tumor epithelium that could potentiate CD8 T cell activity and tumor cell targeting (64–66). Consequently, a blunted MHC class I induction response by tumors cells expressing high levels of NCOR2 would facilitate CD8 T cell immune cell evasion and permit the growth and expansion of disseminated tumor cells (5). Evidence for this paradigm was supported by flow cytometry analysis of cultured 4T07 breast cancer cells treated with IFNγ which revealed a significant increase in MHC class I and B2M expression in the cells in which NCOR2 was knocked down (Figure 6K-M). These data indicate that NCOR2 regulates the lung metastatic outgrowth of breast cancer cells, at least in large part, by repressing tumor cell IFNγ signaling to compromise CD8 T cell activity and killing efficiency through reduced tumor cell expression of MHC class I molecules and compromised apoptosis induction and tumor cell killing.

### NCOR2 modulates lung metastatic outgrowth by regulating IFNγ signaling

We next examined the causal relationship between NCOR2’s ability to repress IFNγ signaling and the level of apoptosis and lung metastatic outgrowth of 4T07 breast cancer cells. We repressed IFNγ signaling in the 4T07 cells with and without NCOR2 knockdown by pre-incubating with a function blocking antibody to the IFNγ receptor and then tested impact on lung metastasis following tail vein injection. Cohorts of syngeneic host mice injected with 4T07 cells expressing firefly luciferase, with and without NCOR2 knockdown, and treated with an IFNγ receptor inhibitory antibody or a matched isotype control were prepared. The mice were monitored twice weekly for metastatic lung growth using bioluminescent imaging and the excised lungs were examined for number, size and phenotype of metastatic lesions at study termination (Figure 7A). We observed a striking effect on metastatic lung outgrowth induced by blocking IFNγ signaling in the breast tumor cells in which NCOR2 had been knocked down (Figure 7B and 7C). While BLI revealed a significant reduction in lung metastases in the mice that had been injected with 4T07 cells in which NCOR2 had been knocked down, those mice that had been also been treated with the anti-IFNγ receptor IgG showed a similar BLI signal as the mice injected with 4T07 cells expressing the control vector (Figure 7B and 7C). H&E analysis of lung tissues clearly showed a significant reduction in macrometastases (overall metastatic burden) and metastatic area in the lungs of the mice injected with 4T07 breast cancer cells in which NCOR2 was knocked down (Figure 7D-G). By contrast, those mice that had been injected with 4T07 tumors in which NCOR2 was knocked down but that had also been incubated with anti-IFNγ receptor function blocking IgG showed a metastatic burden that was remarkably similar to that observed in the mice that had been injected with 4T07 breast cancer cells expressing the control vector (Figures 7D-G). Interestingly, immunohistochemical staining for total leukocytes (CD45+) revealed a higher number of CD45+ cells within the NCOR2-deficient metastatic lung tissue compared to control tissue, but revealed that this difference in infiltrate was lost when IFNγ receptor function was inhibited (Figure 7H and 7I). These data indicate that NCOR2 regulates the metastatic outgrowth of disseminated breast cancer cells by impairing anti-tumor immunity and implicate repression of IFNγ signaling in the tumor cells as a causal factor regulating this phenotype.

**Fig. 7.**
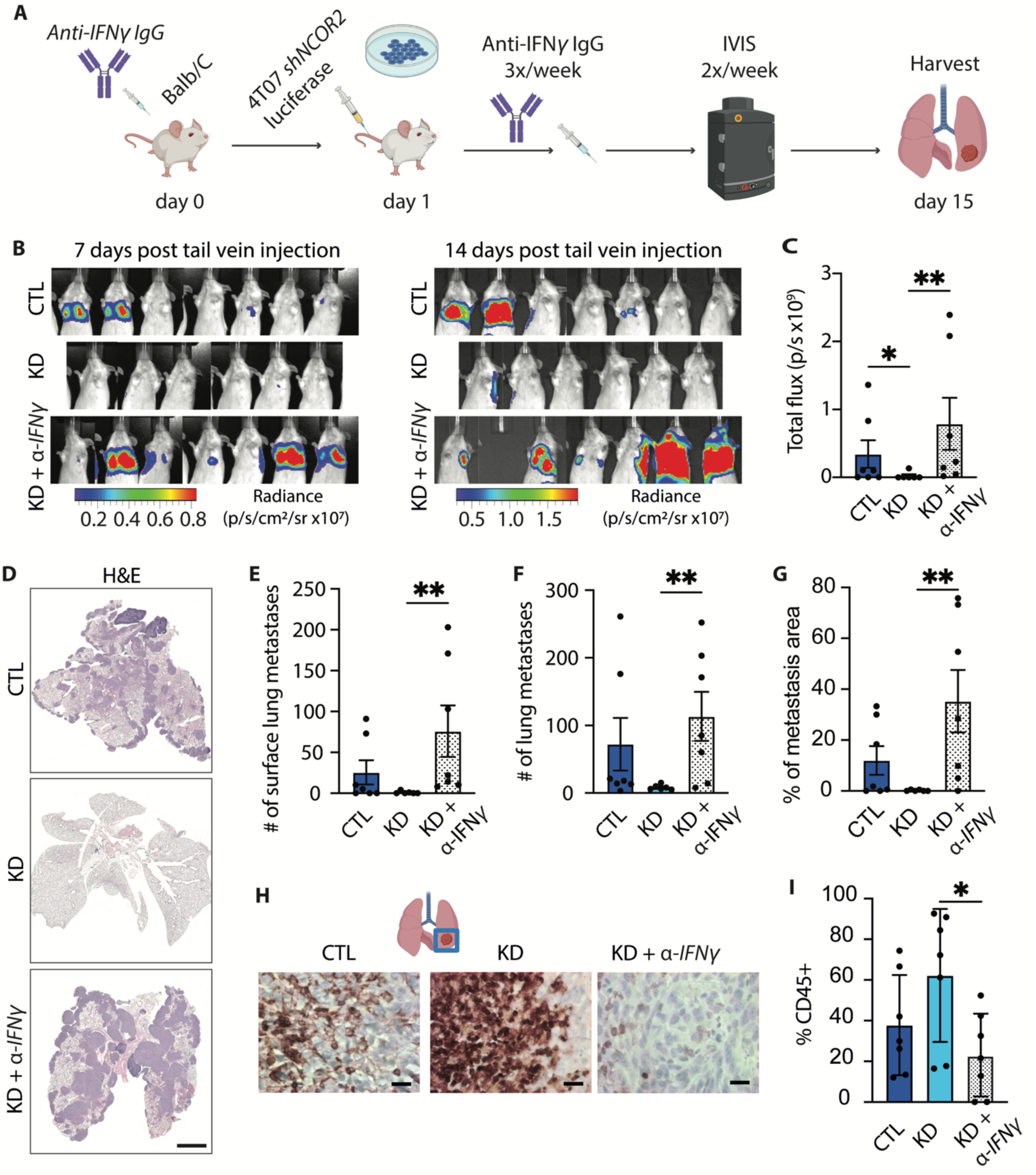
NCOR2 modulates lung metastatic outgrowth through IFNγ signaling. (**A**) Cartoon showing experimental design. (**B**) Representative bioluminescence images of Balb/C mice 7 and 14 days post tail vein injection with 4T07 cells with (KD) and without (CTL) NCOR2 knockdown and antibody to the IFNγ receptor. **(C)** Quantification of Bioluminescence (BLI; total flux) of mouse lung regions from the 14 day time point in (B) (control, *n* = 7; 4T07 NCOR2 knockdown *n* = 6; NCOR2 knockdown and antibody to the IFNγ receptor, *n* = 7). (**D**) Representative images of paraffin lung sections stained for H&E. Scale bar: 1 mm. (**E**) Bar graphs showing quantification of the number of surface lung metastases shown in (D) (control, *n* = 7; NCOR2 knockdown, *n* = 6; NCOR2 knockdown and antibody to the IFNγ receptor, *n* = 7). (**F**) Bar graphs showing quantification of the number of lung metastases shown in (D) (control, *n* = 7; NCOR2 knockdown, *n* = 6; NCOR2 knockdown and antibody to the IFNγ receptor, *n* = 7). (**G**) Bar graphs showing quantification of percent of metastasis area in the lungs shown in (D) (control, *n* = 7; NCOR2 knockdown, *n* = 6; NCOR2 knockdown and antibody to the IFNγ receptor, *n* = 7). (**H**) Representative images of IHC staining for CD45 in the lungs shown in C. Scale bar: 20 μm. (**I**) Bar graphs showing quantification of IHC staining shown in (H) (control, *n* = 7; NCOR2 knockdown, *n* = 7; NCOR2 knockdown and antibody to the IFNγ receptor, *n* = 7). Graphs show mean ± s.e.m. *P < 0.05, **P < 0.01 (two-tailed Mann-Whitney *U* test).

### Association between NCOR2 and MHC class I expression in human breast cancer cells, CD8 T cells and breast cancer metastasis

Our murine studies implicated NCOR2 as an important regulator of the metastatic outgrowth of TNBC cells by repressing IFNγ signaling in the tumor cells to repress CD8 T cell activity and compromise apoptosis induction and MHC class I molecule expression in the DTCs. Accordingly, high expression of NCOR2 in human breast cancers should correlate with reduced MHC class I molecule expression and decreased numbers of activated CD8 T cells in primary and metastatic human breast tumors. Consistent with this prediction, bioinformatics analysis of 10,676 single cells from 6 primary HER2+ and 21,420 single cells from 8 TNBC human tumors that included quantification of NCOR2, HLA-A, HLA-B and B2M mRNA revealed that tumor cells with high NCOR2 expression display a striking inverse correlation with these MHC class I molecules (Figure 8A and 8B). Furthermore, immunohistochemical analysis of primary and lymph node metastatic lesions in human TNBCs revealed a significant inverse correlation between high nuclear NCOR2 expression and CD8 T cell infiltration (Figure 8C-F). These findings provide evidence suggesting that human breast tumors that express high levels of nuclear NCOR2 gain a competitive advantage to form progressive metastatic lesions because they exhibit compromised IFNγ signaling that represses their MHC class I molecule cell surface expression and apoptosis induction in response to T cell secreted IFNγ. Reduced MHC class molecule expression on NCOR2 high tumor cells would also compromise CD8 T cell activation favoring tumor cell survival and metastatic outgrowth (32).

**Fig. 8.**
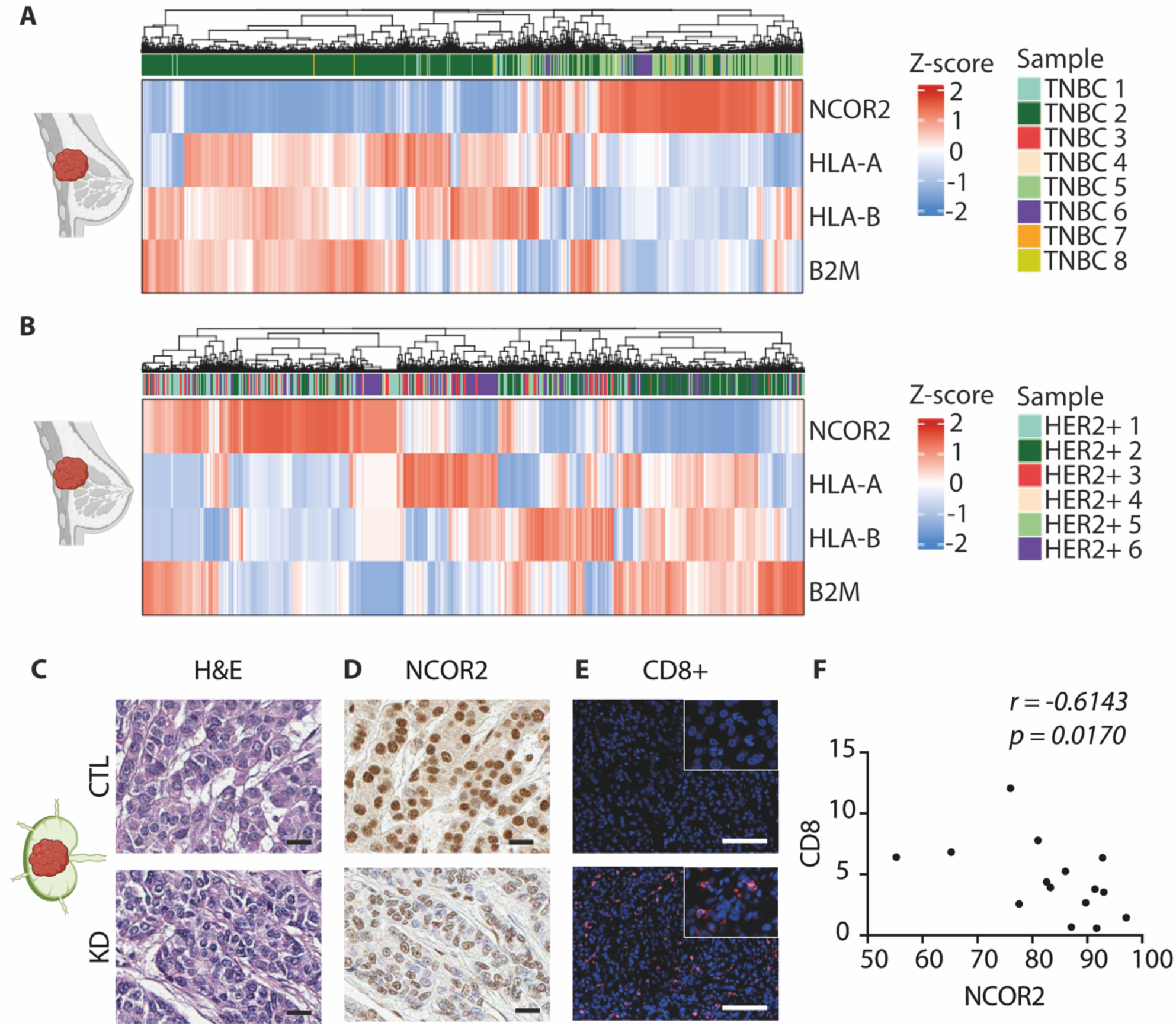
Association between NCOR2, CD8+ T cells, and IFNγ-dependent gene expression in human metastatic breast cancers. (A and. **B)** Heatmap of NCOR2 expression versus MHC class I genes from primary breast cancer patients using publicly available single cell RNA sequencing datasets (See methods), including triple negative (**A**), HER2 positive (**B**). (**C**) Representative images of paraffin sections from human breast cancer lymph node metastasis stained for H&E. Scale bar: 20 μm. (**D**) IHC staining of paraffin sections as in (C) using an antibody against NCOR2. Scale bar: 20 μm. (**E**) IHC staining of paraffin sections as in (C) using an antibody against CD8. Scale bar: 100 μm. (**F**) Scatter plot showing Spierman’s correlation between the percent CD8+ cells and NCOR2+ cells. The correlation coefficient (RS) and P values are shown.

## DISCUSSION

We demonstrate that the nuclear corepressor protein NCOR2 can function as an epigenetic promoter of metastatic progression of breast cancer. Our studies revealed that NCOR2 permits metastasis by repressing tumor cell IFNγ signaling and transcription to prevent the up regulation of MHC class I molecules on tumor cells that permits immune evasion. We showed that reducing tumor cell NCOR2 potentiates IFNγ-dependent killing, permits up regulation of class 1 MHC molecules in disseminated tumor cells and expands the number of activated CD8 positive T cells at the metastatic lung site to restrain metastatic colonization. Consistently, patients whose TNBC and HER2+ breast tumors expressed high levels of NCOR2 also had very low levels of MHC class 1 molecules HLA-A, HLA-B and B2M and the metastatic lesions of patients with TNBC showed a significant inverse correlation between CD8+ T cell infiltration and nuclear NCOR2 expression. The findings expand the repertoire of targetable epigenetic factors known to regulate the transcription of MHC molecules in tumor cells to modulate antitumor immunity (67–70).

Identification of NCOR2 as an epigenetic regulator of MHC class 1 expression presents an attractive antitumor target towards which treatments that impede its function could be designed and implemented to prevent metastatic outgrowth and enhance patient survival.

Immune checkpoint inhibitors have emerged as exciting approaches to leverage the immune system to treat tumors and eradicate metastatic disease (71–75). Although immune checkpoint treatment has shown encouraging promise for some solid cancers, including groups of patients with TNBCs, there remains considerable room for improvement (76–79). A major regulator of CD8+ T cell activity and efficient antitumor immunity is MHC class 1 antigen presentation (35, 38, 39). Not surprisingly, tumors frequently acquire a plethora of irreversible and reversible molecular mechanisms that reduce their MHC molecule expression and/or antigen presentation to thwart antitumor immunity (40, 43, 45). Our findings identify NCOR2 as a key epigenetic repressor of antitumor immunity through its ability to inhibit IFNγ transcription of MHC class 1 molecules. Because NCOR2 can also modulate NF-κB transcription, the discovery expands the repertoire of reversible and targetable mechanisms that compromise IFNγ and NF-κB-signaling tumors adopt to decrease MHC class I molecule expression to evade immune detection and compromise immune therapy (47–51).

Mechanisms dictating dormancy versus metastatic progression include factors regulating the angiogenic switch, stress programs that control tumor cell growth versus stasis, as well as tumor stemness and tumor microenvironmental factors (80–83). Efficient immunosurveillance has emerged as another powerful mechanism dictating dormancy including expression and antigen presentation by class 1 MHC molecules (15–17). We identified NCOR2 as an important promoter of metastatic progression through its ability to modulate IFNγ transcription to regulate expression of MHC class 1 molecules in disseminated tumor cells. Our data also indicate that NCOR2 may suppress a pyroptotic form of cell death in breast cancer cells that is more productive at eliciting anti-tumor immunity. These findings highlight a key role for NCOR2 in dictating early metastatic progression and suggest factors that induce NCOR2 may play a key role in driving tumor recurrence. Consistently, NCOR2 expression is induced by Notch signaling (84, 85), and Notch has been implicated in metastatic recurrence (86, 87). Interestingly, NCOR2 also directly interacts with the dormancy inducing transcription factor orphan hormone receptor NR2F1, and as such may inhibit its function to prevent tumor cell stasis and dormancy to promote metastatic progression (88). Accordingly, clarifying factors regulating NCOR2 levels and function may provide insight into tumor dormancy and metastatic recurrence.

## Acknowledgements

We thank Natashia Korets for mouse colony husbandry and histology support. We thank Dr. M Lazar (University of Pennsylvania) for generously providing the NCOR2 transgenic lines (89). We acknowledge support from UCSF BIDC imaging center. The work was supported by the Breast Cancer Research Foundation grant A132292, the Marks Foundation 20-036-EDV and the National Institute of Health National Cancer Institute R35 CA242447-01A1 to V.M.W., and the Centro Ciencia & Vida, FB210008, Financiamiento Basal para Centros Científicos y Tecnológicos de Excelencia from ANID - Agencia Nacional de Investigación y Desarrollo; Grant 1230021 from FONDECYT - Fondo Nacional de Desarrollo Científico y Tecnológico to H.G.V.

## Author Contributions

V.M.W. Conceptualization and Supervision. K.T. Intellectual input on human breast tumor analysis. Y.Y.C. Provided human TNBC treated and nontreated tissue and pathology analysis. A.I. Mouse tissue pathology. A.D. and J.J.N. TNBC PDX chemotherapy studies.

J.J.N. Transgenic breeding and colony generation. M.Z. Verification of constructs. P.T. and J.J.N. Syngeneic and transgenic study execution and analysis. P.T. Immunostaining and analysis. J.J.N. in vitro IFNγ treatment studies. K.K. and P.T. Immune profiling and analysis. H.G.V. Bioinformatics analysis. P.T., J.J.N. and H.G.V. Figure preparation. V.M.W. manuscript draft. V.M.W., P.T. and J.J.N. Writing-Review. All authors Editing.

## MATERIALS and METHODS

Our research complies with all relevant ethical regulations. The in vivo experiments were performed in accordance with the guidelines from University of California, San Francisco (UCSF). Study protocols were approved by the Institutional Animal Care and Use Committee (IACUC).

### Human breast tissue acquisition and processing

Human breast tumor specimens were obtained from patients with breast cancer from University of California, San Francisco, (UCSF) between 2010 and 2020. Tissue specimens were flash frozen in OCT (Tissue-Tek) by slow immersion in liquid nitrogen or placement on dry ice and kept at −80°C until cryosectioning. All specimens were de-identified, stored, and analyzed according to the procedures described in institutional review board (IRB) protocol #10-03832, approved by the UCSF Committee of Human Resources IRB (Pro00034242). Human breast cancer tissue microarrays were obtained from US Biomax (Rockville, MD, USA).

### Cell culture

4T1 cells and 4T07 cells were grown on tissue culture plastic in RPMI with 10% FCS and antibiotics, HEK-293 cells were grown on tissue culture plastic in DMEM with 10% FBS and antibiotics. Growth medium was changed 1 day after plating and every 2 days thereafter. Cell lines were tested for Mycoplasma (MycoAlert Mycoplasma Detection Kit; Lonza). All cell lines were derived from authenticated sources.

### Cell growth curve

4T07 cells were grown on tissue culture plastic in RPMI with 10% FCS and antibiotics. Part of the cells was pretreated with 2 μL/mL of 0.1 IPTG for 4 days. Thereafter, the cells were seeded at a density of 7000 cells per well in a corning 12 well plate (3.6 cm^2^, Fisher Scientific). Counting was carried out using hemocytometer chamber each 24 hours for 6 days.

### Lentiviral infection and vectors

Stable knockdown of NCOR2 was achieved by lentivirus-mediated RNA interference (RNAi). Sequences of validated shRNA against murine NCOR2 were obtained from the Broad Institute Genetic Perturbation Platform (<u>GPP Web Portal - Gene Details</u> (broadinstitute.org) and cloned into two custom built 2nd generation (shRNA) IPTG inducible vectors (Supplemental figure 1A) between AgeI and EcoRI sites using the U6 inducible 3 x LacO developed at the Broad Institute (<u>Vector Details: pLI_913</u> (broadinstitute.org). Murine NCOR2 shRNA included TRCN0000238139 (TGTTGGCTTACAGGGTATATT) validated in this study, and TRCN000095283 (CCAGATCATCTACGATGAGAA), and TRCN000095281 (CCAGTGTAAGAACTTCTACTT) validated by Sigma Mission. A shRNA targeting EGFP was used as non-specific control (GCAAGCTGACCCTGAAGTTCAT) 4T1 and 4T07 cells transduced with these constructs were selected with up to 1 mg/ml G418 and hairpins induced with 200 µM IPTG (*in vitro*) or 20 mM Dioxane Free IPTG in drinking water provided ad libidum (*in vivo*).

### Transgenic experiments

NCOR2 knockout mice (PyMT MMTV-Cre NCOR2 KO) were generated by crossing MMTV-PyMT with MMTV-Cre NCOR2 KO mice. Animal husbandry for mice was carried out in Laboratory Animal Resource Center (LARC) facilities at UCSF. Once a week, tumors were measured using calipers. Mice were sacrificed at 13-22 week once a single tumor reached 1.5cm in diameter. Mice were anesthetized with 5% inhalant isoflurane. Then, under anesthesia, retroorbital blood was collected from the venous sinus and a transcardial perfusion was performed by opening the thoracic cavity. A blunt tip needle was inserted in the right ventricle and a small incision was made in the left atrium. Transcardial perfusion was performed with 30–40 mL PBS. While still attached in situ, lungs were then directly perfused via the trachea with 1mL 4% formaldehyde in order to avoid collapsing. The lungs were dissected, surface metastases were macroscopically calculated, and the tissues were paraffin-embedded for subsequent analysis. Mammary tumors from euthanized mice were harvested and paraffin-embedded, cryo-embedded, and snap frozen tumor chunks were collected.

### Tail vein experiments

For tail vein experiments, 4T07 cells with or without NCOR2 knockdown and luciferase expression were used. 4 days before the injection, cells were pretreated with IPTG. Female Balb/C and NSG mice were obtained from The Jackson Laboratory. Animal husbandry for mice was carried out in Laboratory Animal Resource Center (LARC) facilities at UCSF. 8 weeks old female Balb/C mice were administrated IPTG in drinking water. Following day, mice were injected with 10^6^ cells suspended in 100 μl of PBS into the tail vein. Lung metastases were monitored biweekly by bioluminescence imaging. Administration of IPTG continued until sacrifice 14 days following tail vein injection. 29 weeks old female Balb/C mice were administrated IPTG water 5mg/mL. Following day, mice were injected with 10^6^ cells suspended in 100 μl of PBS into the tail vein. Lung metastases were monitored biweekly by bioluminescence imaging. Administration of IPTG continued until sacrifice 10 days following tail vein injection. 9 weeks old female Balb/C mice were administrated IPTG water 5mg/mL. Then, mice were divided into three groups. First group was injected intraperitoneally with 500 ug control IgG (Rat IgG1 Isotype Control, Leinco Technologies) diluted in PBS; following day, mice were injected with 10^6^ 4T07 control cells suspended in 100 μl of PBS into the tail vein. Second group was injected intraperitoneally with 500 μg control IgG (Rat IgG1 Isotype Control, Leinco Technologies) diluted in PBS; following day, mice were injected with 10^6^ 4T07 NCOR2 knockdown cells suspended in 100 μl of PBS into the tail vein. Third group was injected intraperitoneally with 500 ug anti-IFNγ (anti-Mouse IFNγ, Leinco Technologies) diluted in PBS; following day, mice were injected with 10^6^ 4T07 NCOR2 knockdown cells suspended in 100 μl of PBS into the tail vein. Lung metastases were monitored biweekly by bioluminescence imaging. Administration of IPTG continued until sacrifice 14 days following tail vein injection. At the end of experiment, lung perfusion and animal euthanasia were performed as described above and lungs were paraffin-embedded. For flow cytometry analysis, lungs were perfused only with PBS and stored in a mixture of 95% FBS 5% DMSO at -80 °C.

### Orthotopic experiments

For orthotopic experiments, 4T1 and 4T07 cells with or without NCOR2 knockdown were used. Cells were pretreated with 2 μL/mL of 0.1 IPTG 4 days before the injection. Female Balb/C mice were obtained from The Jackson Laboratory. Animal husbandry for mice was carried out in the Laboratory Animal Resource Center (LARC) facilities at UCSF. 16 weeks old female Balb/C mice were administrated IPTG water 5mg/mL. Following day, mice were anesthetized using 3.5% inhalant isoflurane and Buprenorphine and Meloxicam intraperitoneal anesthesia and ophthalmic ointment was applied. Hair over the area surrounding left 4^th^ mammary gland were removed and the skin was wiped using 70% alcohol followed by 2% chlorhexidine solution. Then, 0.5cm incision between the 4^th^ nipple and the midline was made and skin was gently separated from the underlying fascia using a cotton swab and blunt dissection. 4^th^ fat pad was located and stabilized using micro-dissecting forceps and injected with a mixture of 10^5^ cells 4T1 cells with or without NCOR2 knockdown suspended in 50 μl of PBS and 50 of Matrigel. Then, the incision was closed, and topical local analgesic (Lidocaine) was applied. Postsurgical intraperitoneal anesthesia with Buprenorphine and Meloxicam was performed. Tumors were measured using caliper biweekly. Administration of IPTG continued until sacrifice 4 weeks following mammary fat pad injection. At the end of experiment, retroorbital blood collection, lung perfusion and animal euthanasia were performed as described above. The lungs were dissected, surface metastases were macroscopically calculated, and the tissues were paraffin-embedded for subsequent analysis. Mammary tumors from euthanized mice were harvested and paraffin-embedded, cryo-embedded, and snap frozen tumor chunks were collected. 7 weeks old female Balb/C mice were orthotopically injected as described above with 4T07 cells with or without NCOR2 knockdown. Mice were sacrificed 4.5 weeks following mammary fat pad injection and the tissues were harvested as described above.

### Bioluminescence imaging

Bioluminescence imaging was performed using the IVIS Imaging System (Xenogen; Caliper Life Sciences) within the IACUC animal barrier at UCSF. Mice were anesthetized with 2% isoflurane and injected intraperitoneally with 150 mg/kg D-luciferin 10 min before image acquisition. Lung metastatic tissue was determined by measuring the photon flux from a region of interest drawn around the bioluminescence signal using the Live Imaging Software v.4.5.2 (PerkinElmer).

### Histology, Image Acquisition, and Analysis

Prior to staining, paraffin-embedded tissue sections were dewaxed and hydrated through a descending grade ethanol series. Then sections were stained with Mayer’s hematoxylin solution (Sigma Aldrich) for 8 min and with Eosin Y solution (Sigma Aldrich) for 30 s. The tissue sections were mounted with Cytoseal (Richard Allen Scientific). Images were acquired digitally using Leica Biosystems Microscope Slide Scanner Aperio AT2 and visualized using ImageScope software. Quantification of number of lung metastases was performed by manually. To score percentage of metastasis area within the whole lung, percentage of metastatic area per total lung area was calculated using QuPath.

### Immunohistochemistry, Image Acquisition and Analysis

Prior to staining, paraffin-embedded tissue sections were dewaxed and hydrated through a descending grade ethanol series. After microwave treatment for 10 minutes in tris-based antigen unmasking solution (Vector Laboratories), endogenous peroxidase activity was blocked with 3% H2O2 for 10 minutes. Tissues sections were then blocked with 5% bovine serum, albumin 5% goat serum in PBS for 1 h at room temperature. Sections were then incubated overnight at 4°C with primary antibodies specific to CD8a (Invitrogen, clone SP16), CD45 (BD Pharmingen, clone 30-F11: 1:200), cleaved caspase-3 (Cell Signaling, catalog: 1:200), Ki-67 (Invitrogen, catalog: 1:200) and NCOR2 (Sigma Aldrich). Thereafter, sections were incubated with Impress anti-rabbit or anti-rat reagent (Novus Biologicals) for 30 min at room temperature. For diaminobenzidine (DAB) staining, positive signal was developed with Impress DAB reagent (Vector Laboratories). Sections were counterstained with hematoxylin and mounted with Cytoseal. Alternatively, tyramide signal amplification (TSA) was applied instead of DAB staining: sections were incubated with (Alexa Fluor, Thermo Fisher Scientific: 1:500) diluted in Tyramide Amplification Buffer (Biotium) for 10 min. Cell nuclei were then counterstained with DAPI, sections were mounted with Vectashied, and imaged. Images of DAB-stained sections were acquired using an Olympus IX81 microscope at 40x magnification with a 0.6 Ph2 objective. Images were acquired with a Photometrics CoolSNAP HQ2 CCD camera using Infinity Capture v6.5.6 software. Full-slide images were acquired digitally using Leica Biosystems Microscope Slide Scanner Aperio AT2 and visualized using ImageScope software. Quantification of positive staining was performed by manually counting nuclei of interest and total nuclei and taking an average or using QuPath. At least 5 images of each tissue specimen were used for averaged analyses.

Images of TSA-stained sections were acquired using a Nikon Eclipse TE2000-U inverted microscope at 20x magnification with a 0.75 NA objective. Images were acquired with a Hamamatsu ORCAFlash4.0 LT camera using NIS-Elements software. Full-slide images were acquired digitally using Zeiss Axio Scan Z.1. Quantification of tissue sections was performed by thresholding images at a fixed intensity and counting the percentage of positive cell nuclei per total nuclei using Fiji. 5 images of each tissue specimen were used for averaged analyses.

### Immunofluorescence, Image Acquisition and Analysis

Frozen OCT tissue 10 μm sections were fixed in 4% PFA for 10 min at RT, and then permeabilized with 0.25% v/v Triton-X-100 in PBS for 5 min at room temperature. Tissues sections were then blocked with 5% bovine serum, albumin 5% goat serum in PBS for 1 h at room temperature. Primary antibody specific to NCOR2 (Sigma Aldrich) was diluted in 5% bovine serum in PBS and incubated overnight at 4°C. Tissue sections were then stained with fluorescently labelled secondary antibodies (Alexa Fluor, Thermo Fisher Scientific: 1:500) diluted in 5% bovine serum albumin in PBS for 1 h at room temperature. Tissue sections were counterstained with DAPI, mounted with Vectashied, and imaged. Images of IF-stained sections were acquired using a Nikon Eclipse TE2000-U inverted microscope at 20x magnification with a 0.75 NA objective. Images were acquired with a Hamamatsu ORCAFlash4.0 LT camera using NIS-Elements software. Quantification of tissue sections was performed by thresholding images at a fixed intensity and counting the percentage of positive cell nuclei per total nuclei using Fiji. 10 images of each tissue specimen were used for averaged analyses.

### Flow cytometry

Mouse lung tissue was thawed and chopped with a razor blade. Chopped tissue was digested in 100 U ml−1 collagenase type 1 (Worthington Biochemical, catalogue #: number LS004196), 500 U ml−1 collagenase type 4 (Worthington Biochemical, catalogue number LS004188) and 200 µg ml−1 DNase I (Roche, catalogue number 10104159001) in DMEM while shaking at 37 °C. Digested tissue was filtered using a 100 µm filter to remove remaining pieces. Red blood cells were lysed in ACK buffer (Thermo Fisher Scientific, catalogue number A1049201) and remaining cells were counted. Samples were then washed with FACS buffer (2% FBS in PBS) and resuspended in appropriate buffer for staining for flow cytometric analysis.

For flow cytometric analyses, cells were washed with PBS prior to staining with Zombie NIR Fixable live/dead dye (Biolegend) for 20 min at 4°C. Cells were washed in PBS followed by surface staining for 30 min at 4°C with directly conjugated antibodies diluted in FACS buffer containing anti-CD16/32 (clone 2.4G2; BioXCell) to block non-specific binding. Cells were washed again with FACS buffer prior to read-out on a BD LSR Fortessa SORP cytometer. The following antibodies were used: anti-CD25 FITC (Biolegend, dilution 1:200, clone 3C7), anti-CD11b APC (eBioscience, dilution 1:100, clone M1/70), anti-CD90.2 AF700 (Biolegend, dilution 1:400, clone 30-H12), anti-CD8 PerCp-Cy7 (Biolegend, dilution 1:400, clone 53-6.7), anti-CD45 BV421 (eBioscience, dilution 1:400, clone 30-F11), anti-CD45R BV785 (Biolegend, dilution 1:400, clone RA3-6B2), anti-CD4 BUV395 (Biolegend, dilution 1:400, clone RM4-5), anti-CD44 BUV737 (Biolegend, dilution 1:400, clone IM7)

### Flow cytometry for cell death, MHC class I and β2-microglobulin cell surface expression

4T07 cells expressing IPTG-inducible NCOR2-targeting shRNAs or GFP-targeting shRNA were cultured as described and treated with IPTG for 72 hrs prior to the reapplication of IPTG (1 mM) and stimulation with and without IFNγ for 48 hrs. For quantification of cell death, cell media was collected and attached cells were harvested by trypsinization for staining with the Live-or-Dye NucFix™ Red Staining Kit (Biotium; Cat: #: 32010-T) for 30 mins prior to two washes with PBS and fixation in 2% paraformaldehyde (PFA). For assays examining cell surface expression of MHC class I molecules and β2-microglobulin, cells were treated in the same manner as above but also co-treated with the caspase inhibitors Ac-DEVD-CHO and Ac-IETD-CHO for 24 hrs prior to and throughout the duration of the experiment to avoid loss of cells due to apoptosis. Following treatment, cell media was collected and attached cells were harvested by trypsinization for blocking in PBS+2% Fetal bovine serum (FBS; FACS wash buffer), mouse serum and TruStain FcX™ (anti-mouse CD16/32) Antibody (BioLegend; Cat.#: 101320) for 30 mins followed by staining with a PE anti-mouse H-2Kd/H-2Dd Antibody (MHC Class I, clone 34-1-2S, BioLegend; Cat. #: 114708) and a PE/Cyanine7 anti-mouse β2-microglobulin Antibody (clone A16041A, Cat. #:154508) for an additional 30 minutes. DAPI (4’,6-diamidino-2-phenylindole) was then added for 10 mins to distinguish live/dead cells before washing cells with a large volume of FACS wash buffer and fixing cells in 2% PFA. Flow cytometry was performed using a BD LSRFortessa™ Cell Analyzer, and population percentages were defined and quantified using FlowJo™ v10.8 Software (BD Life Sciences).

### RNA extraction and RT-qPCR-based detection of circulating tumor cells

Total RNA was extracted from blood and purified using the RNeasy Mini Kit (QIAGEN). Total RNA (1.0 μg) was used as a template for cDNA synthesis using M-MLV reverse transcriptase (Promega Corporation). cDNA (100 ng) was used as template for PCR amplification using the LightCycler FastStart DNA MasterPLUS SYBR Green I Kit (Roche) and the LightCycler System (Roche). Oligonucleotide primers were designed using the LightCycler Probe Design Software v.2.0 (Roche). Transcript expression was quantified by normalizing the gene of interest copy number (per μl) to absolute levels of an endogenous, stably expressed reference gene HPRT. Analysis of the data was performer using 2-ΔΔCT method described previously (90, 91).

### Western blotting

Protein lysates were prepared using radioimmunoprecipitation assay (RIPA; 50 mM Tris-HCl, 150 mM NaCl, 0.25% Na-deoxycholate, 0.1% SDS, and 1.0% IGEPAL CA-630 [NP-40]; pH7.4 at room temperature) or Laemmli lysis buffer (50 mM Tris-HCl, 2% SDS, and 5 mM EDTA; pH 7.4 at room temperature). Immediately before cell lysis, a cocktail of protease inhibitors (1.2 μg leupeptin, 1.2 μg pepstatin, 2.4 μgaprotinin, 12 μg E-64, 0.5 mM benzamidine, 50 mM NaF, and 1.2μg Pefabloc) and the phosphatase inhibitor Na-orthovanadate (1 mM activated with 1.5% H2O2) were added to the buffer. RIPA lysates were carried out on ice, while Laemmli lysates were performed at room temperature due to SDS precipitation at cold temperatures. After rinsing the dishes with PBS, lysis buffer was added, and cells were scraped off the dish with a cell scraper. RIPA lysates were sonicated three times, each for 10s, while Laemmli lysates were passed through a fine pipette tip (p200) several times. Both were centrifuged for 10 min at 20,817 relative centrifugal force to pellet any internal organelles and cellular debris. Supernatant was collected and fast frozen on dry ice. Protein concentration was determined using a bicinchoninic acid assay kit (Promega) following the manufacturer’s protocol. Protein extracts were obtained with a kit using the manufacturer’s instructions (Thermo Fisher Scientific). Equal amounts of protein were separated on reducing SDS-PA gels, immuno-blotted, and detected with the ECL Plus System. Samples were boiled for 5 min (95°C) and loaded onto the SDS-PA gel, and protein was separated at 120 constant volts. The protein was transferred onto a prewet polyvinylidene difluoride membrane (100% methanol, 1 min) at 300 mA for 2 h. The polyvinylidene difluoride membrane was rinsed with Tris-buffered saline with Tween 20 (TBST), and nonspecific binding was blocked with 5% nonfat dry milk dissolved in TBST. The membrane was then incubated with the primary antibody overnight at 4°C, washed with TBST, incubated with HRP-conjugated secondary anti-body (1h; room temperature; dilution, 1:2000), washed with TBST, and detected with the chemiluminescence system Quantum HRP substrate (Advansta). Signal quantification and normalization to loading control was performed using Fiji.

### Single cell RNAseq analyses of primary breast cancer tumors

We downloaded the single cell RNAseq data of primary (92) from Gene Expression Omnibus (https://www.ncbi.nlm.nih.gov/ geo/) using series GSE161529. The Seurat pipeline was applied to each sample (93). Raw counts of annotated cancer cells were normalized (Seurat NormalizeData function) and scaled (ScaleData function. The expression of selected genes then visualized in heatmaps using the R package ComplexHeatmap (94).

### Statistical analysis

GraphPad Prism Version 10.0.2 was used to perform all statistical analyses and correlations and statistical significance was determined using the appropriate tests as noted in the corresponding figure legends. Tests of normality that incorporated skewness and kurtosis were used to determine the appropriate statistical test. All independent variables are described in the figure legends with measurements always from distinct samples (biological replicates). All tests are two-tailed.

