## Extended data for "NCOR2 represses MHC class I molecule expression to drive metastatic progression of breast cancer"

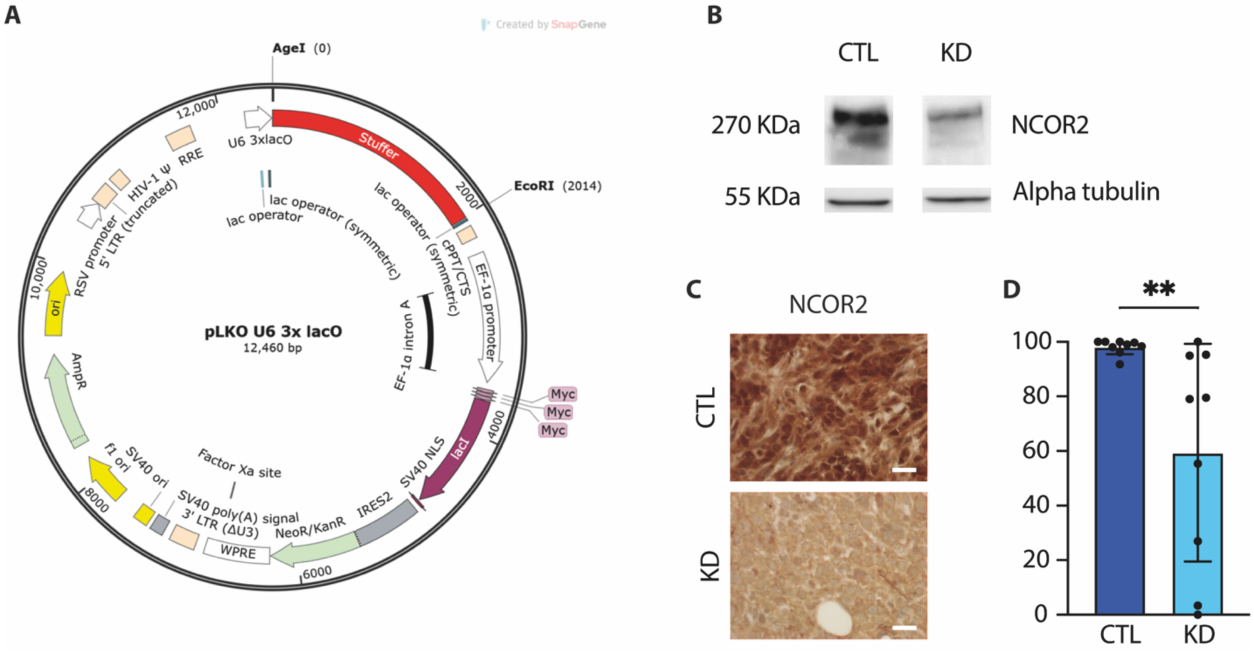


**Extended Data Fig. 1. NCOR2 knockdown validation in 4T1 mammary tumors. (A)** Plasmid map illustrating the cloning vector used to induce breast cancer cells with luciferase cDNA for bioluminescence imaging and an inducible shRNA sequence targeting either *GFP* (CTL) or *NCOR2* (KD). **(B)** Representative immunoblots of NCOR2 showing shRNA-mediated knockdown of NCOR2 expression in 4T1 cells (KD) as compared to a GFP-targeting shRNA (CTL), with alpha tubulin used as a loading control. (**C**) Representative images of immunohistochemical (IHC) staining for NCOR2 in paraffin sections from 4T1 syngeneic primary tumors. Scale bar: 20 µm. (**D**) Bar graphs showing quantification of IHC NCOR2 staining shown in C (control, $n$ = 9; NCOR2 knockdown, $n$ = 9). Graphs show mean ± s.e.m. **P < 0.01 (two-tailed Mann-Whitney $U$ test).

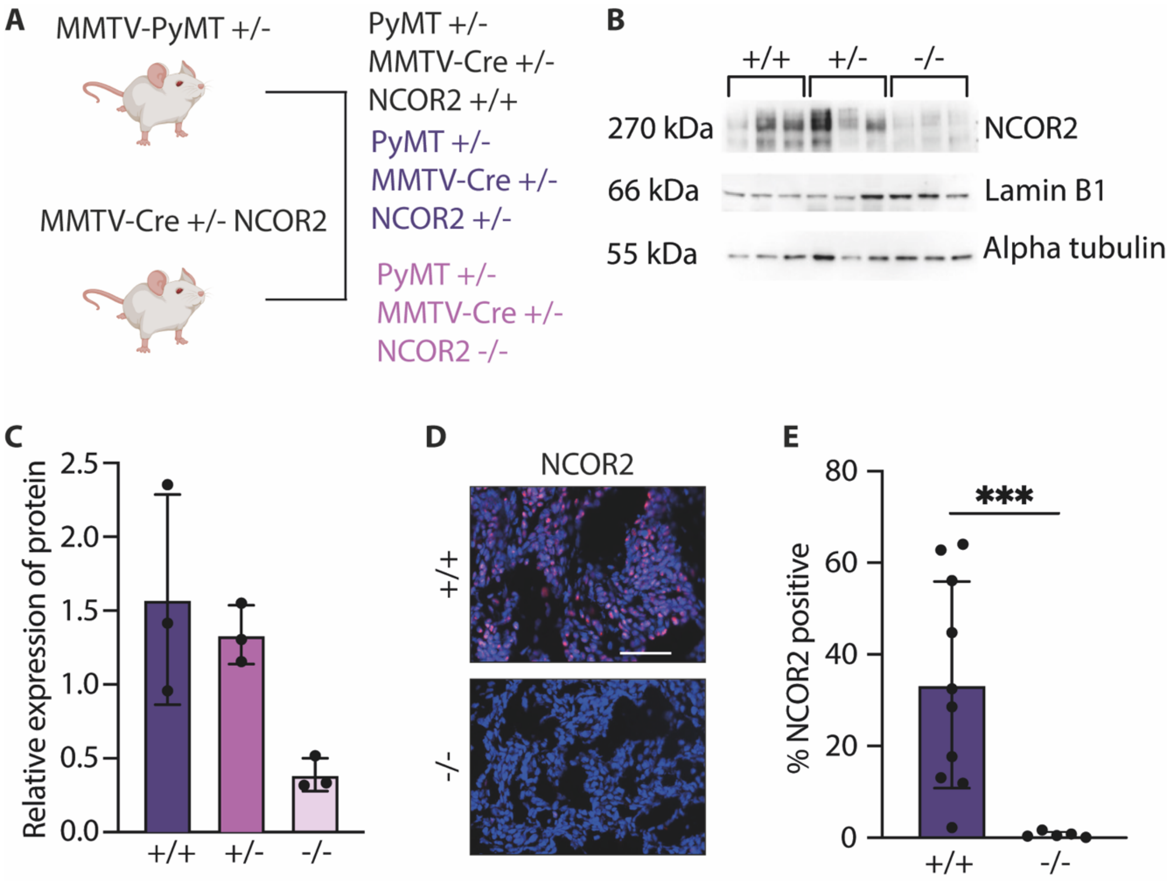


**Extended Data Fig. 2. NCOR2 knockout validation in spontaneous PyMT-driven mammary tumors. (A)** Cartoon illustrating breeding strategy to generate genotypes with wildtype NCOR2 (+/+), and heterozygous (+/-) and homozygous (-/-) knockout of NCOR2. (**B**) Representative immunoblots of transgenic primary tumors showing knockout of NCOR2, with lamin B1 and alpha tubulin used as a loading control. (**C**) Quantification of relative expression of NCOR2 for B (PyMT wild type (+/+), $n = 3$; PyMT NCOR2$^{+/-}$, $n = 3$; PyMT NCOR2$^{-/-}$, $n = 3$). (**D**) Representative images of immunohistochemical (IHC) staining for NCOR2 in frozen sections of primary transgenic tumors. Scale bar: 50 μm. (**E**) Bar graphs showing quantification of positive IHC staining shown in C (PyMT wild type (+/+), $n = 10$; PyMT NCOR2$^{-/-}$, $n = 5$). Graphs show mean ± s.e.m. ***$P < 0.001$ (two-tailed Mann-Whitney $U$ test).

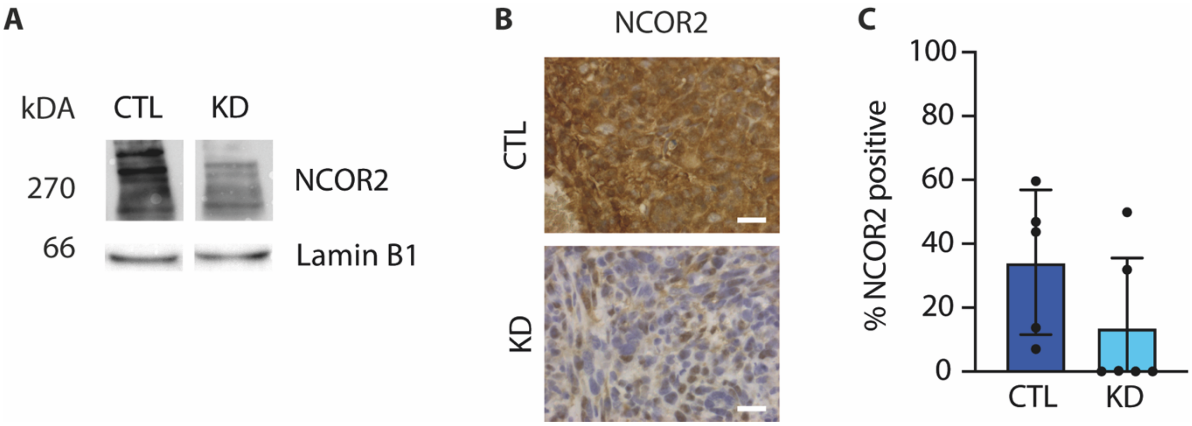


**Extended Data Fig. 3. NCOR2 knockdown validation in 4TO7 mammary tumors. (A)** Representative immunoblots of NCOR2 showing shRNA-mediated knockdown of NCOR2 expression in 4T07 cells (KD) as compared to a GFP-targeting shRNA (CTL), with lamin B1 as a loading control. (**B**) immunohistochemical (IHC) staining for NCOR2 in paraffin sections of lungs harvested 14 days after tail vein injection with 4T07 cells with or without knockdown of NCOR2. Scale bar: 20 µm. (**C**) Bar graphs showing quantification of IHC staining shown in B (control, *n* = 5; NCOR2 knockdown, *n* = 6). Graph shows mean ± s.e.m.

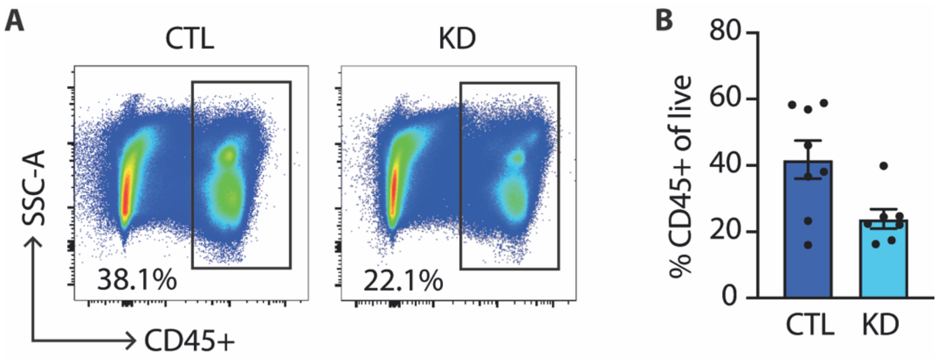


**Extended Data Fig. 4. Percentage of CD45-positive cells in the lungs of mice receiving intravenous injection of 4TO7 cells with or without NCOR2 knockdown. (A)** Representative flow cytometry plots showing the fraction of CD45+ cells isolated from the lungs of syngeneic mice from Figure 6. **(B)** Bar graphs showing quantification of the percentage of CD45+ cells, calculated from total live cells (control, $n$ = 8; NCOR2 knockdown, $n$ = 7). Graphs show mean ± s.e.m.

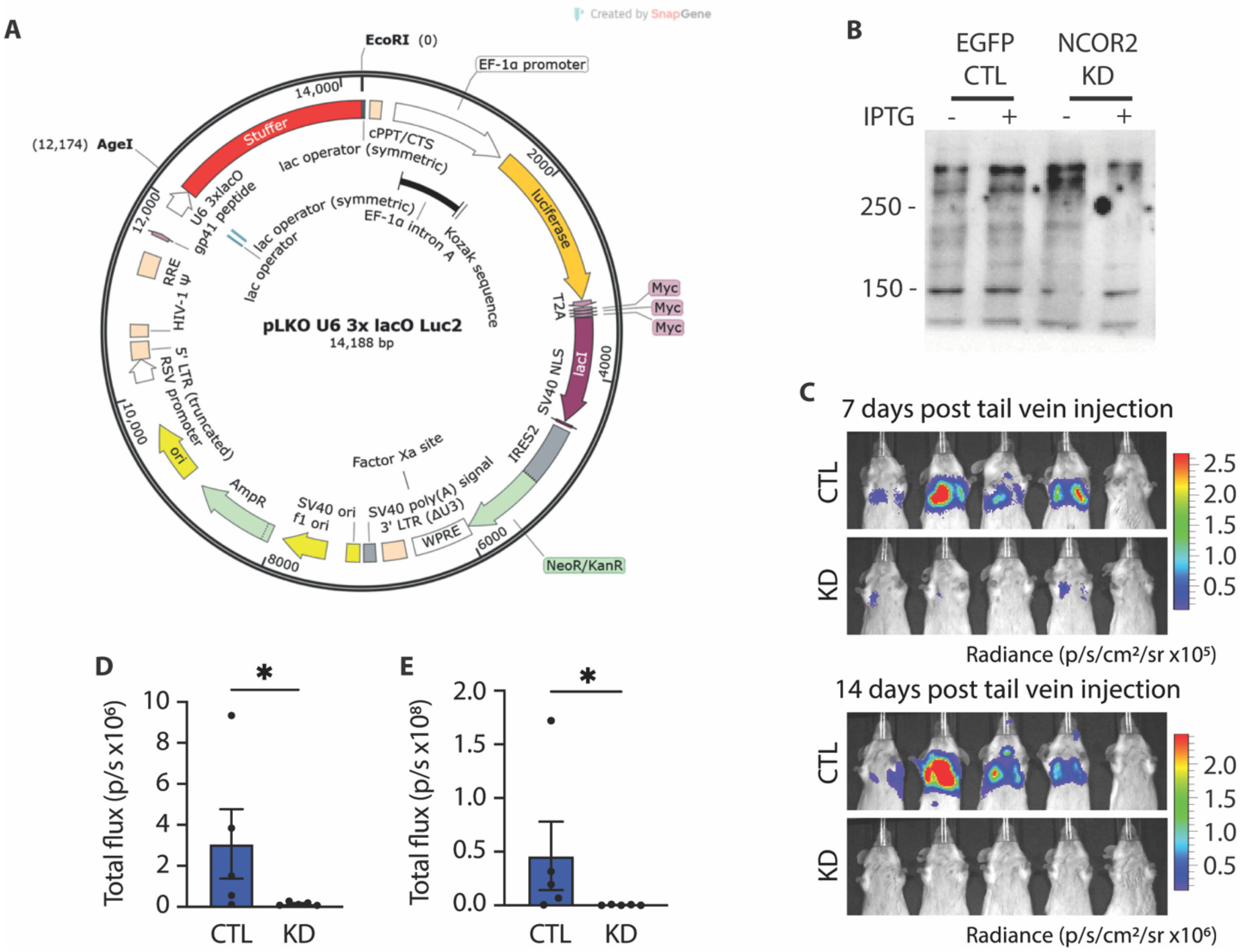


**Extended Data Fig. 5. Validation of knockdown and reduced experimental lung metastasis using a second NCOR2-targeting shRNA in 4TO7 mammary tumor cells. (A)** Plasmid map illustrating the newer version of the cloning vector used to induce breast cancer cells with luciferase cDNA for bioluminescence imaging and an inducible shRNA sequence targeting either *GFP* (CTL) or *NCOR2* (KD). **(B)** Representative immunoblots of NCOR2 showing shRNA-mediated knockdown of NCOR2 expression in 4T07 cells (KD) as compared to a GFP-targeting shRNA (CTL). Ponceau staining was used to confirm equal loading of samples. **(C)** Representative bioluminescence images of Balb/C mice 7 days and 14 days after tail vein injection with 4T07 cells with or without knockdown of NCOR2. **(D and E)** Quantification of total flux (photons/second) for CTL and KD mice from (C) at 7 days (D) and 14 days (E) post tail vein injection (control, CTL; *n* = 9; NCOR2 knockdown, KD; *n* = 9). Graphs show mean ± s.e.m. *P < 0.01 (two-tailed Mann-Whitney *U* test).
